# Hybrid histidine kinase activation by cyclic di-GMP-mediated domain liberation

**DOI:** 10.1101/675454

**Authors:** Badri N. Dubey, Elia Agustoni, Raphael Böhm, Andreas Kaczmarczyk, Francesca Mangia, Christoph von Arx, Urs Jenal, Sebastian Hiller, Iván Plaza-Menacho, Tilman Schirmer

## Abstract

Cytosolic hybrid histidine kinases (HHKs) constitute major signalling nodes that control various biological processes, but their input signals and how these are processed are largely unknown. In *Caulobacter crescentus*, the HHK ShkA is essential for accurate timing of the G1-S cell cycle transition and is regulated by the corresponding increase in the level of the second messenger c-di-GMP. Here, we use a combination of X-ray crystallography, NMR spectroscopy, functional analyses and kinetic modelling to reveal the regulatory mechanism of ShkA. In the absence of c-di-GMP, ShkA predominantly adopts a compact domain arrangement that is catalytically inactive. C-di-GMP binds to the dedicated pseudo-receiver domain Rec1 thereby liberating the canonical Rec2 domain from its central position where it obstructs the large-scale motions required for catalysis. Thus, c-di-GMP cannot only stabilize domain interactions, but also engage in domain dissociation to allosterically control activity. Enzyme kinetics data are consistent with conformational selection of the ensemble of active domain constellations by the ligand.

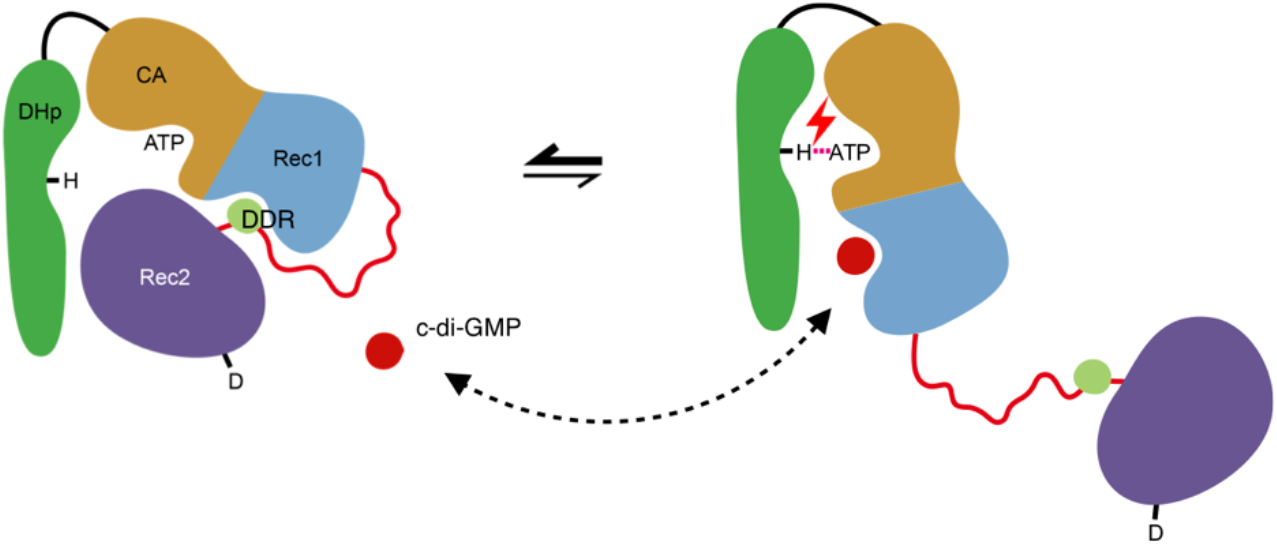

## Introduction

Two-component systems (TCSs) are widespread in microorganisms, and mediate signal-dependent regulation of gene expression. They link diverse intracellular and extracellular stimuli to specific cellular responses including development, cell division, and antibiotic resistance^1,2^. In their simplest form, TCSs are composed of a sensory histidine kinase (HK) and its cognate response regulator. The sensory HK is usually a homodimer that auto-phosphorylates a histidine in response to an input signal and then catalyzes transfer of the phosphoryl group to a conserved aspartate on the receiver domain (Rec) of its partner response regulator, thereby modifying its activity. As well as a Rec domain, response regulators usually harbour an output effector domain that often functions as a transcription factor. The HK catalytic core is composed of a dimerization and histidine transfer (DHp) domain for carrying the active histidine, and a catalytic ATP binding (CA) domain. The DHp domain contains two α-helices that mediate homodimerization by forming a four-helix bundle. An exposed histidine residue in the DHp domain is autophosphorylated by the CA domain, which forms an α/β-sandwich that binds ATP^3^. Phosphorelays, also called multicomponent systems, are more complex in that they include a histidine phosphotransferase (Hpt) domain to enable phosphate transfer between two Rec domains. In hybrid histidine kinases (HHK) that function as part of phosphorelays, the first Rec domain is fused C-terminally to the histidine kinase core. Although HHKs comprise almost 20 % of bacterial histidine kinases ^4^, no detailed structural or mechanistic studies on their regulation are available.

Typically, HKs perceive the signal by dedicated periplasmic domains, but also membrane spanning and cytosolic domains/proteins are known to regulate them^5^. Input domains, such as PAS, GAF and HAMP, are located N-terminal to the HK core and affect the conformation and dynamics of the dimeric DHp helix bundle upon signal recognition, thereby controlling auto-phosphorylation of the conserved histidine on DHp α1^6–8^ and initiating signal transduction. In contrast, ShkA, investigated here, is a non-canonical HHK lacking any N-terminal input domain, but exhibiting a pseudo-receiver domain (Rec1) between the kinase core and the C-terminal receiver domain (Fig. 1a). In the accompanying study^9^, it is shown that such domain organization is common and that the pseudoreceiver domain provides the binding site for the input signal.

**Figure 1.**
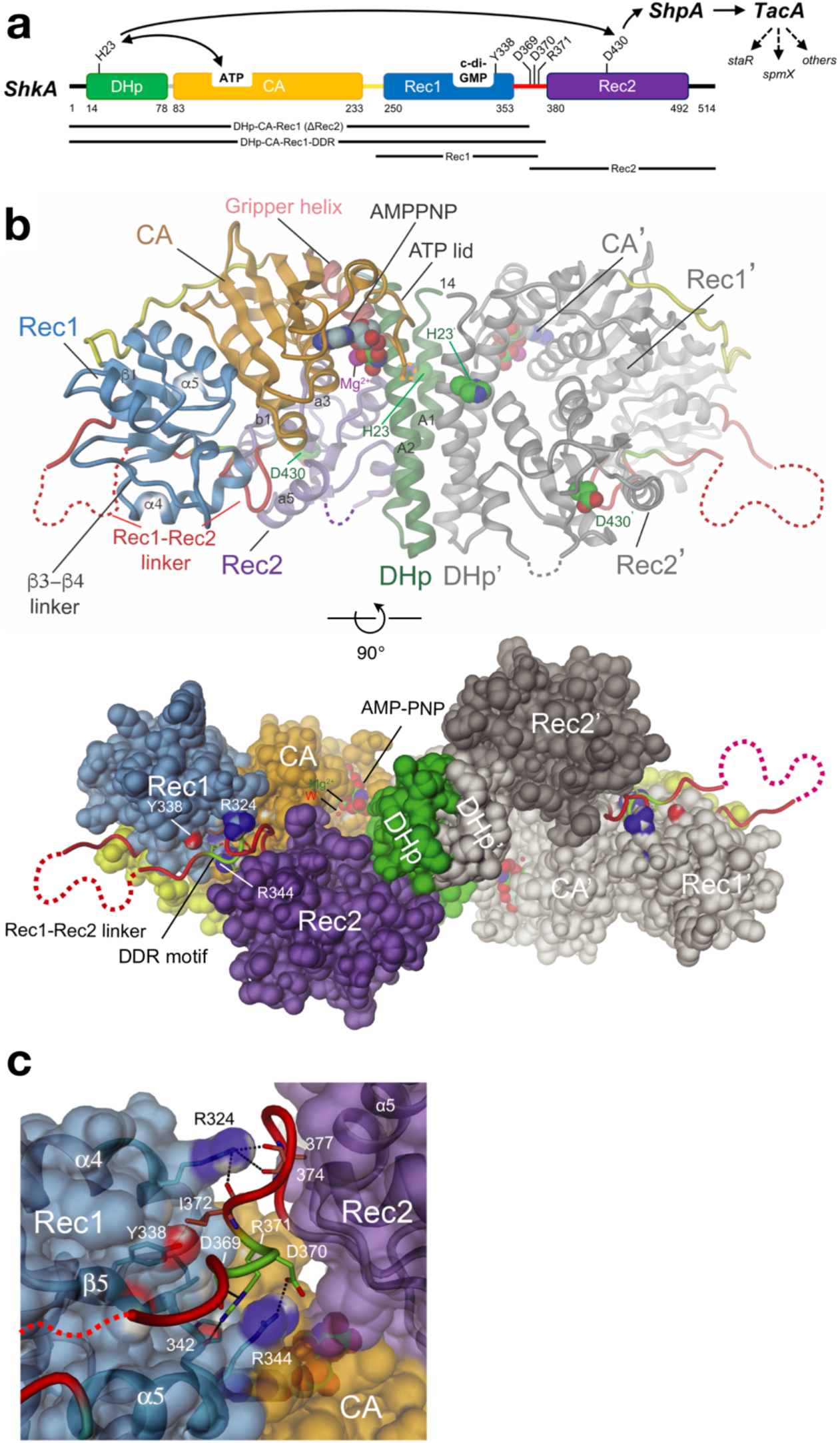
Crystal structure of full-length ShkA dimer shows compact, auto-inhibited domain arrangement. **(a)** ShkA domain organisation with functional residues indicated. The phospho-relay from ShkA *via* ShpA to transcription factor TacA, which controls the expression of *staR, spmX*, and other genes (broken arrows), is indicated by solid arrows. **(b)** Side and bottom view of SkhA dimer in cartoon or surface representation with one protomer colored as in panel A and the other (primed labels) in gray. Disordered loops are indicated by dashed lines. His23, Asp430, and bound AMPPNP/Mg^++^ are shown in CPK representation. The structure represents an inactive, auto-inhibited conformation, since these potentially reacting groups are dispersed. For more structural details, see **Supplementary Fig. 1**. **(c)** Detailed view of Rec1-Rec2 linker (red, with DDR motif residues in light-green) bound to the α4 - β5 - α5 face of Rec1. Residues of the motif interact with Rec1 main-chain carbonyl 342 and R344 side-chain. Main-chain carbonyls 374 and 377 of the linker interact with Rec1 R324, and I372 forms an apolar contact (with I327). Despite the absence of direct Rec1/Rec2 contacts, these interactions tether tightly the N-terminus of Rec2 to Rec1.

The second messenger c-di-GMP has been recognized as a major player in bacterial signal transduction, regulating diverse functions including bacterial life style, behavior, and development^10^. c-di-GMP is produced by diguanylate cyclases, and diguanylate cyclase domains (GGDEF), together with phosphodiesterase (EAL) domains^11^, comprise the second most common group of TCS output domains, establishing a firm link between the two major systems in bacterial signaling. A connection in the opposite direction has been discovered only recently in *Caulobacter crescentus* with the cell cycle histidine kinase CckA being controlled by c-di-GMP^12^. Increasing levels of c-di-GMP during the G1 phase cause CckA to switch from default kinase to phosphatase mode upon S phase entry, thereby licensing cells for replication initiation. Based on structural and functional analyses, c-di-GMP mediated cross-linking of the DHp and CA domain was identified as the molecular mechanism of the switch^13^. Most recently it was found that ShkA, the other major kinase involved in the regulation of the *C. crescentus* cell cycle progression and morphogenesis^14^, is also controlled by c-di-GMP^9^. However, in contrast to CckA, which upon binding of c-di-GMP switches into phosphatase mode, the ligand is required for ShkA kinase activity suggesting a distinct mechanism of allosteric control.

Here, we reveal the molecular mechanism of ShkA regulation, which is based on c-di-GMP mediated mobilization of a locked, auto-inhibited domain arrangement. We have determined crystal structures of full-length ShkA and of its isolated Rec1 domain in complex with c-di-GMP and have analyzed the dynamics of the full-length enzyme by NMR spectroscopy employing isoleucine methyl group isotope labelling. Comprehensive enzymatic analyses revealed the underlying thermodynamics of the regulatory mechanism. Finally, a general model for the structural transitions during the catalytic cycle of HHKs is proposed.

## Results

### Full-length ShkA crystal structure reveals an auto-inhibited state of ShkA

ShkA is a hybrid histidine kinase composed of a DHp-CA core domain, which catalyzes autophosphorylation of H23, and a Rec2 receiver domain carrying the phospho-acceptor D430 (Fig. 1a). A second, but degenerate, receiver domain Rec1 precedes Rec2. The crystal structure of full-length ShkA was determined by molecular replacement to 2.8 Å resolution in the presence of AMPPNP/Mg^2+^ (Table 1). The structure shows a crystallographic dimer stabilized by contacts mediated by the DHp helices A1 and A2 (Fig. 1b, Supplemental Fig. 1a). The DHp domain is followed by a canonical CA domain with a tightly associated Rec1 domain (Supplemental Fig. 1b). Rec1 shows the canonical (βα)5 receiver domain fold (see also Fig. 3a), but with the fourth cross-over helix degenerated to a loop (β3 - β4 loop). The acidic pocket is incomplete in that it has the two canonical acidic groups at the end of β1 replaced by S255, P256, explaining why the domain is not involved in phosphotransfer, despite the presence of an aspartate (D297) in the canonical phosphoacceptor position. Finally, the partly disordered Rec1-Rec2 linker leads to the canonical Rec2 domain that is found associated with the central DHp bundle (Supplemental Figs. 1c, d). Notably, the N-terminal end of the Rec2 domain appears firmly tethered to Rec1 not through a direct contact but *via* multiple ionic interactions mediated mainly by a DDR motif on the Rec1 - Rec2 linker (light green, in Figs. 1b, c).

**Table 1.**
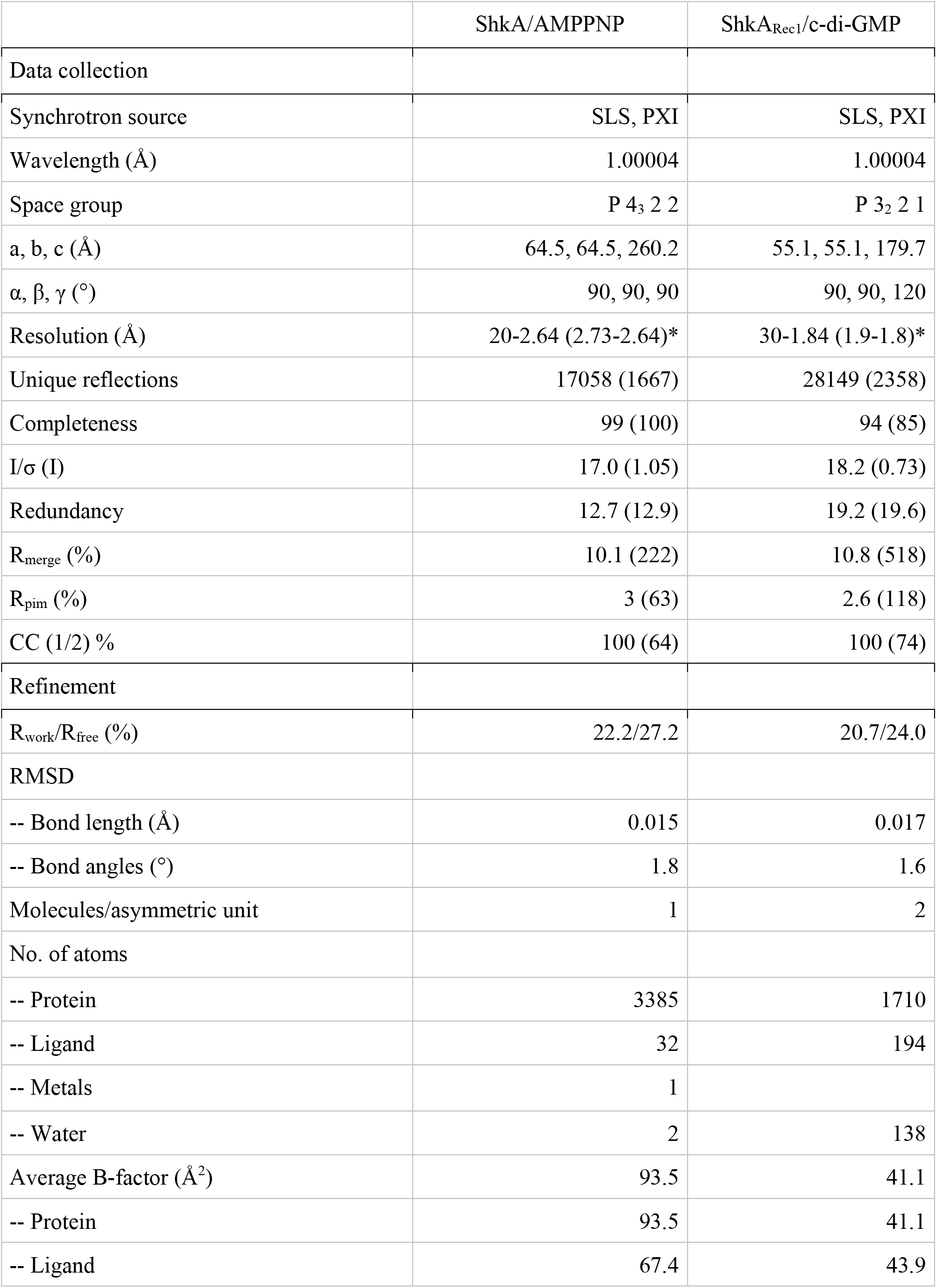
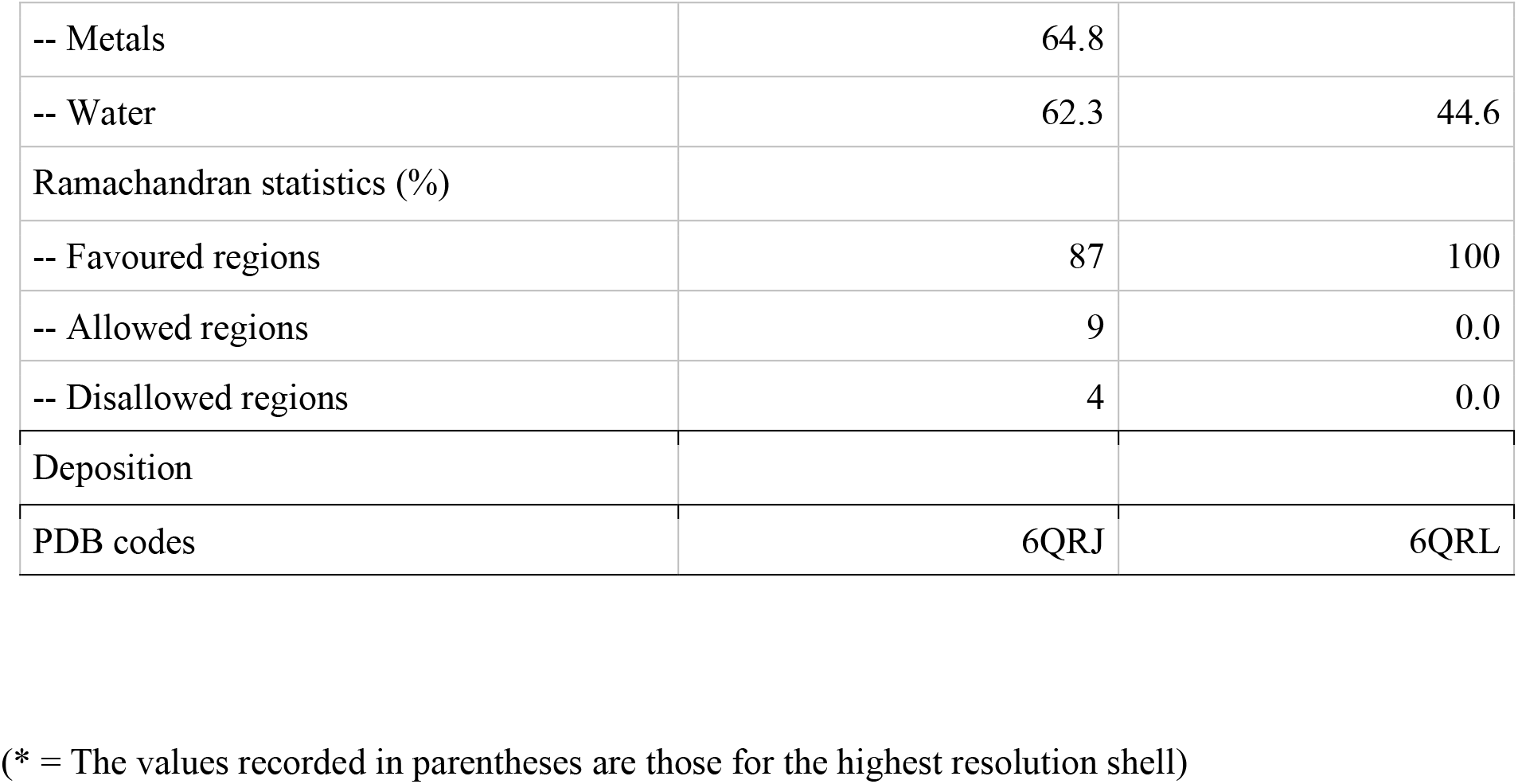
Crystallographic data collection and refinement statistics.

|  | ShkA/AMPPNP | ShkA<sub>Rec1</sub>/c-di-GMP |
| --- | --- | --- |
| Data collection |  |  |
| Synchrotron source | SLS, PXI | SLS, PXI |
| Wavelength (Å) | 1.00004 | 1.00004 |
| Space group | P 4<sub>3</sub> 2 2 | P 3<sub>2</sub> 2 1 |
| a, b, c (Å) | 64.5, 64.5, 260.2 | 55.1, 55.1, 179.7 |
| α, β, γ (°) | 90, 90, 90 | 90, 90, 120 |
| Resolution (Å) | 20-2.64 (2.73-2.64)* | 30-1.84 (1.9-1.8)* |
| Unique reflections | 17058 (1667) | 28149 (2358) |
| Completeness | 99 (100) | 94 (85) |
| I/σ (I) | 17.0 (1.05) | 18.2 (0.73) |
| Redundancy | 12.7 (12.9) | 19.2 (19.6) |
| R<sub>merge</sub> (%) | 10.1 (222) | 10.8 (518) |
| R<sub>pim</sub> (%) | 3 (63) | 2.6 (118) |
| CC (1/2) % | 100 (64) | 100 (74) |
| Refinement |  |  |
| R<sub>work</sub>/R<sub>free</sub> (%) | 22.2/27.2 | 20.7/24.0 |
| RMSD |  |  |
| -- Bond length (Å) | 0.015 | 0.017 |
| -- Bond angles (°) | 1.8 | 1.6 |
| Molecules/asymmetric unit | 1 | 2 |
| No. of atoms |  |  |
| -- Protein | 3385 | 1710 |
| -- Ligand | 32 | 194 |
| -- Metals | 1 |  |
| -- Water | 2 | 138 |
| Average B-factor (Å<sup>2</sup>) | 93.5 | 41.1 |
| -- Protein | 93.5 | 41.1 |
| -- Ligand | 67.4 | 43.9 |
| -- Metals | 64.8 |  |
| -- Water | 62.3 | 44.6 |
| Ramachandran statistics (%) |  |  |
| -- Favoured regions | 87 | 100 |
| -- Allowed regions | 9 | 0.0 |
| -- Disallowed regions | 4 | 0.0 |
| Deposition |  |  |
| PDB codes | 6QRJ | 6QRL |

(\* = The values recorded in parentheses are those for the highest resolution shell)

The structure clearly represents an inactive, auto-inhibited, state of ShkA. Large conformational changes would be required to allow access of bound ATP to H23 and of H23~P to D430 for autophosphorylation and phosphotransfer, respectively. Similarly, DHp mediated dephosphorylation of D430~P would require a substantial rotation of Rec2 towards the putative hydrolytic water thought to reside close to the active histidine^15^.

### C-di-GMP binding induces large conformational and dynamic changes in ShkA

C-di-GMP binds to the Rec1 domain as shown by NMR studies with the isolated domain^9^. To investigate the mechanism of c-di-GMP-mediated relief of the auto-inhibited, locked state (Fig. 1b), we next investigated the full-length enzyme by NMR. A 2D [^15^N,^1^H]-TROSY spectra of uniformly deuterated ShkA and ShkA/AMPPNP show only around 40 strong signals in the random-coil region of the spectrum (Fig. 2a, Supplemental Fig. 2a), indicating that they arise from residues located in locally flexible regions of the protein. The large majority of the 489 non-proline residues of ShkA are however not detected in this spectrum, due to the large molecular size of the rigid dimer and the associated slow molecular tumbling.

**Figure 2.**
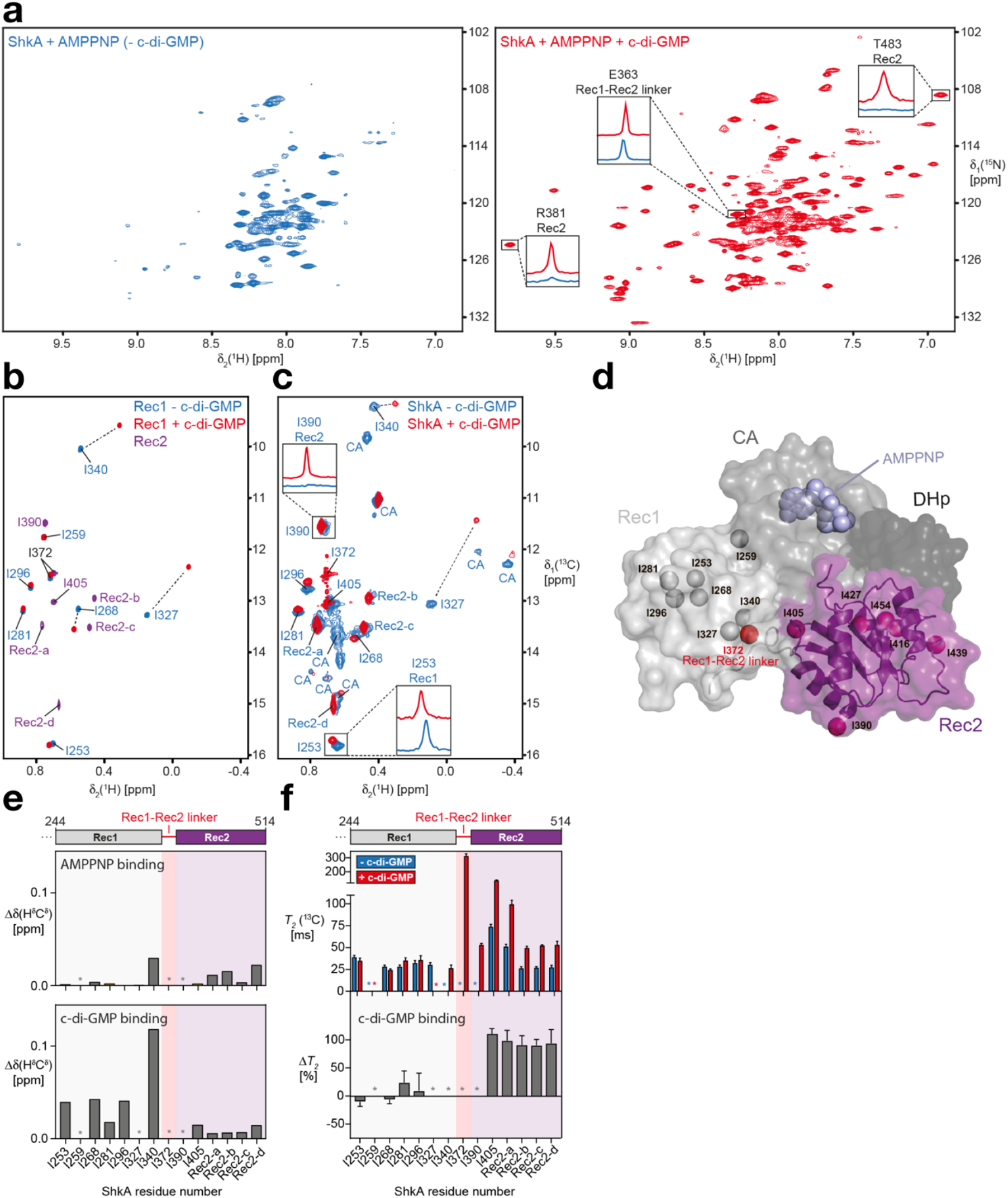
C-di-GMP binding to ShkA leads to release of the Rec2 domain. **(a)** 2D [^15^N,^1^H]-TROSY spectra of ShkA/AMPPNP (blue spectrum) and ShkA/AMPPNP/c-di-GMP (red spectrum). The insets show 1D [^1^H]-cross sections for representative amide protons. **(b)** Overlay of 2D [^13^C,^1^H]-HMQC spectra of the isolated [^13^C/^1^H^δ1^-Ile]Rec1 domain in apo state (blue) and c-di-GMP-bound state (red), and the isolated [^13^C/^1^H^δ1^-Ile]Rec2 domain (purple). **(c)** Spectral overlay of 2D [^13^C,^1^H]-HMQC spectra of [^13^C/^1^H^δ1^-Ile]ShkA in apo state (blue) and c-di-GMP-bound state (red). The two insets show 1D [^1^H]-cross sections for representative methyl protons. Sequence-specific resonance assignments of the δ1 methyl groups are either indicated or assigned to the Rec2 or CA domain. **(d)** Crystal structure of ShkA/AMPPNP. Surface representation of the monomer composed of four domains: DHp (dark grey), CA with bound AMPPNP (grey), Rec1 (light grey) and Rec2 (purple). The δ1 methyl groups of isoleucine residues in the Rec1 and Rec2 domains are shown as spheres together with I372 (red sphere) that is part of the Rec1 – Rec2 linker. **(e)** Chemical shift perturbation of isoleucine δ1 methyl groups upon addition of AMPPNP to ShkA (top) and upon addition of c-di-GMP to ShkA/AMPPNP (bottom). **(f)** Transverse relaxation times of isoleucine δ1 methyl carbons of ShkA/AMPPNP (blue bars) and ShkA/AMPPNP/c-di-GMP (red bars). The bottom panel shows the relative change of *T_2_*(^13^C) upon addition of c-di-GMP (*ΔT_2_*). **(e,f)** Asterisks indicate methyl groups that are either not assigned or line-broadened beyond detection in at least one of the two states.

Binding of AMPPNP to ShkA does not lead to major changes in the spectrum. In contrast, binding of c-di-GMP to either apo ShkA or ShkA/AMPPNP leads to dramatic spectral changes with around 120 well-resolved additional resonances being clearly detected, indicating a substantial change in the dynamics of ShkA for a large part of the protein (Fig. 2a and Supplemental Fig. 2a). This massive increase in signal intensity is exemplified for selected signals with their 1D ^1^H-cross sections (Fig. 2a and Supplemental Fig. 2d). A spectral overlay of ShkA/c-di-GMP with the isolated Rec2 domain shows that the new resonances perfectly overlap (Supplemental Fig. 2a), including Rec2 residues that are in contact with either the CA or DHp domain in the ShkA crystal structure (A421, G422, R446, A447, K448, A471, A472, G473). This indicates that upon binding of c-di-GMP to full-length ShkA, the Rec2 domain loses its contacts with the other domains and detaches completely from the otherwise rigid main architecture of the protein (Supplemental Fig. 2a). Since no significant chemical shift differences are detected between the two spectra, the detached Rec2 domain in ShkA has essentially the same three-dimensional structure as the isolated Rec2 domain.

In order to obtain site-specific information of the effect of c-di-GMP on the structure and dynamics of ShkA, we increased the experimental sensitivity by employing methyl-NMR spectroscopy of specifically δ1-[^13^C/^1^H]-isoleucine labelled full-length ShkA and isolated Rec1 and Rec2 domains (Figs. 2b-d, Supplemental Fig. 2b). Sequence-specific resonance assignments for Rec1 were obtained by identifying unambiguous NOEs in agreement with the crystal structure. Additional assignments for isoleucine methyl groups δ1-I340, δ1-I390 and δ1-I405 were obtained using selective mutagenesis of full-length ShkA (Supplemental Fig. 2c). The mutation I259V located within the Rec1 domain caused major spectral changes, indicating the importance of residue I259 in stabilizing the Rec1–CA domain interface (Supplemental Fig. 1b).

AMPPNP binding to the CA domain of full-length ShkA leads to only minimal chemical shift perturbation of the δ1-Ile methyl groups of the entire protein, implying that the structures of ligand-free and AMPPNP-bound ShkA are very similar (Fig. 2e and Supplemental Fig. 2b). In contrast, binding of c-di-GMP to either apo ShkA or to ShkA/AMPPNP causes substantial spectral changes. All δ1-Ile methyl group signals of the Rec2 domain strongly increase in intensity (Supplemental Fig. 2c), in full agreement with the spectral behavior of the backbone amides. Similar chemical shift changes were observed for c-di-GMP binding either to the isolated ShkA-Rec1 domain or to fulllength ShkA, showing that c-di-GMP binds to the β5-α5 surface of prototypical REC domains (Figs. 2c, e and Supplemental Fig. 2b).

Overall, these data indicate that c-di-GMP binding to ShkA leads to liberation of the Rec2 domain from the protein core. To validate this hypothesis, we conducted NMR spin relaxation experiments in the presence and absence of c-di-GMP (Fig. 2f and Supplemental Fig. 2f). For the isolated receiver domains, average ^13^C transverse relaxation times for ILV methyl groups of Rec2 and Rec1 are quite similar (174 ms and 160 ms, respectively), in agreement with the similar molecular weight of both domains (16.3 kDa and 12.5 kDa, respectively). For full-length ShkA/AMPPNP in the absence of c-di-GMP, the transverse relaxation times of all δ1-Ile resonances is smaller (25 – 75 ms), as expected from the respective protein sizes (Fig. 2f). Addition of c-di-GMP strongly increases the ^13^C *T*_2_ times of Rec2 such that ^13^C *T*_2_ times are effectively doubled, but does not significantly affect those of δ1-Ile of Rec1 (Fig. 2f and Supplemental Fig. 2f). These experiments thus quantitatively confirm that c-di-GMP binding to ShkA leads to the specific detachment of the Rec2 domain from the protein core into a dynamic, multi-conformational state, which enables the enzyme to undergo the large motions required for the phosphoryl transfer reactions.

### C-di-GMP competes with Rec1-Rec2 linker for binding to Rec1 domain

To reveal the molecular mechanism of c-di-GMP-induced Rec2 domain mobilization, we set out to determine the structure of the ShkA/c-di-GMP complex. Well-diffracting crystals were obtained for the isolated Rec1 domain, but not for full-length ShkA/c-di-GMP, and the structure was determined to 1.8 Å resolution (Figs. 3a, b, Table 1). The asymmetric unit contains two monomers each with a bound monomeric c-di-GMP ligand, which are virtually identical and superimpose closely with the corresponding domain in full-length ShkA (Supplement Fig. 3b). In addition, an intercalated c-di-GMP dimer is observed that mediates a 2-fold symmetry contact between the two Rec1 monomers that is most likely a crystal artefact. Indeed, the ITC data of full-length ShkA are consistent with 1:1 stoichiometry (Supplemental Fig. 4).

**Figure 3.**
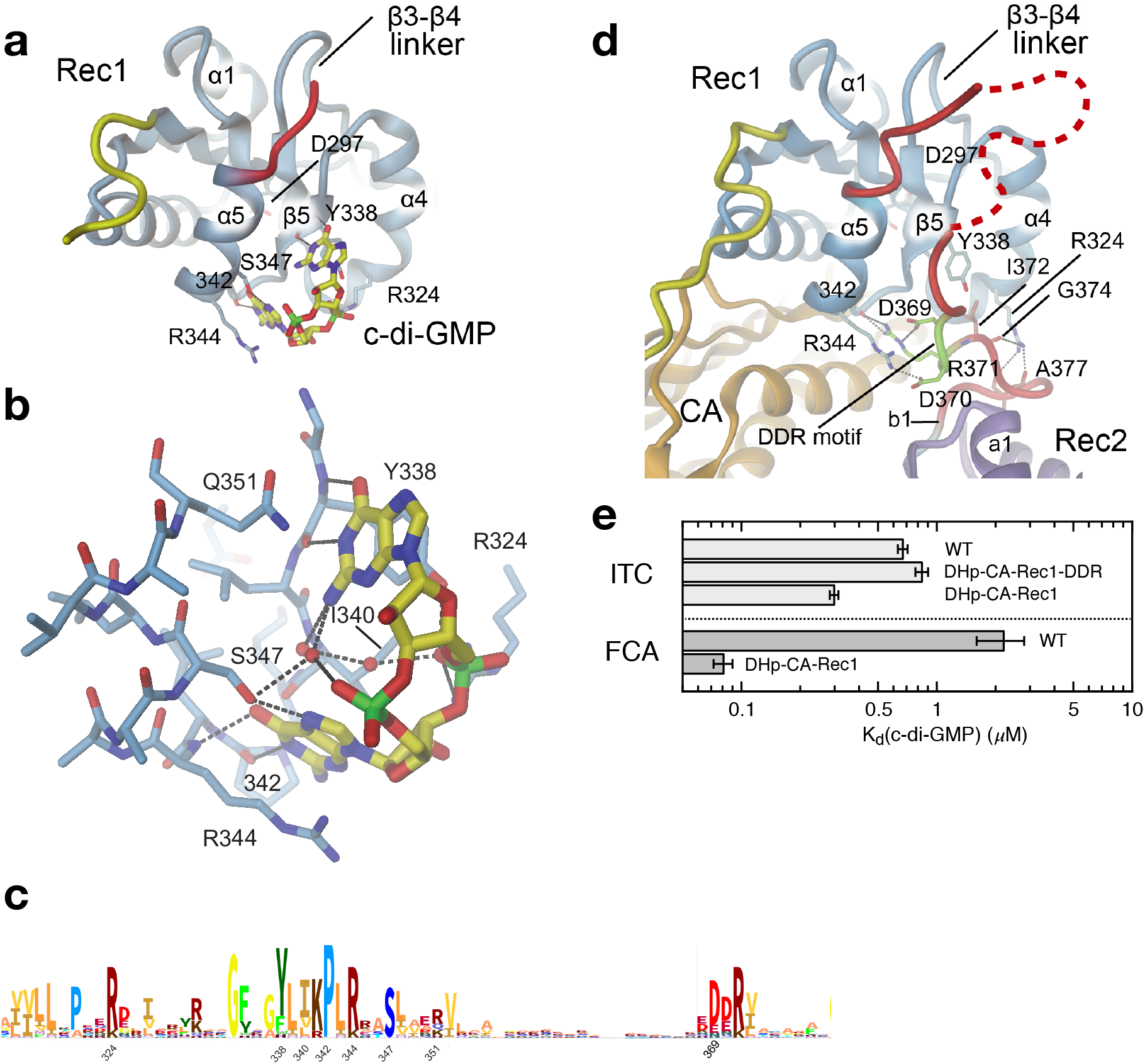
C-di-GMP and Rec1-Rec2 linker compete for Rec1 binding. **(a, b)** Crystal structure of ShkA_Rec1_ in complex with c-di-GMP with interacting residues shown in full. The Watson-Crick edges of both guanines form H-bonds with Rec1 main-chain atoms. The bases stack with Y338, Q351 and with R344. The hydroxyl of S347 is H-bonded to a guanine N7 group. Both Y338 and R324 interact with one of the phosphates. See also **Supplementary Fig. 3a.** **(c)** Sequence logo encompassing the c-di-GMP binding site of the ShkA_Rec1_ domain. **(d)** Structure of Rec1 of full-length ShkA in same orientation as ShkA_Rec1_ in panel a. Comparison with (a) shows that the DDR motif (light-green) is bound to the same site on Rec1 as c-di-GMP, for a superposition see **Supplementary Fig. 3b**. Note that R324 and R344 of Rec1 are involved both in c-di-GMP and linker binding. **(e)** C-di-GMP binding to various ShkA constructs as measured by ITC (top) and fluorescence competition assay (FCA, bottom). Construct DHp-CA-Rec1 shows a significantly lower dissociation constant compared to the other variants, which can be attributed to the absence of the competing DDR segment. Full data are given in **Supplementary Fig. 4.**

Monomeric c-di-GMP is bound to the α4-β5-α5 face of Rec1 (Figs. 3a,b, Supplemental Fig. 3a), i.e. to the surface that is involved in the dimerization of canonical Rec domains. The Watson-Crick edges of both guanine bases form H-bonds with the backbone of the exposed edge of the β5 edge and the β5-α5 loop. In addition, Y338 and R344 are forming stacking and cation - π interactions with the guanyl bases, respectively. S347 is H-bonded to a guanyl base, and R324 forms an ionic interaction with a c-di-GMP phosphate. All interacting ShkA residues are highly conserved in related sequences (Fig. 3c).

In full-length ShkA, the c-di-GMP binding site is occupied by the Rec1-Rec2 linker (Figs. 1c, 3d). Specifically, residues shown to be engaged in c-di-GMP binding (Y338, R324, and R344) are interacting with a conserved DDR motif of the linker region. Thus, we reasoned that if the DDR segment were to compete with c-di-GMP binding, its deletion should increase c-di-GMP affinity. Indeed, affinity measurements by ITC and fluorescence competition showed that the DHp-CA-Rec1 construct had a lower dissociation constant compared to the other investigated variants (Fig. 3e). Why this difference is considerably more pronounced in the fluorescence data is not clear.

Taking together all data on structure and dynamics, we propose a “ligand-coupled conformational equilibrium” (LCCE) model as the mechanism underlying c-di-GMP mediated ShkA activation (Figs. 4a, b). In the absence of c-di-GMP, the enzyme is in a dynamic equilibrium between a closed, autoinhibited state (Ec) and an ensemble of open, inhibition-relieved states (Eo) that are characterized by a liberated Rec2 domain. In absence of the ligand, this equilibrium would be populated largely on the Ec side, in line with the NMR results and the constitutive inactivity of the enzyme. C-di-GMP mediated ShkA activation would then proceed by conformational selection^16^ of the Eo states with their unobstructed Rec1 binding site, shifting the equilibrium in a dose-dependent fashion to the active states.

**Figure 4.**
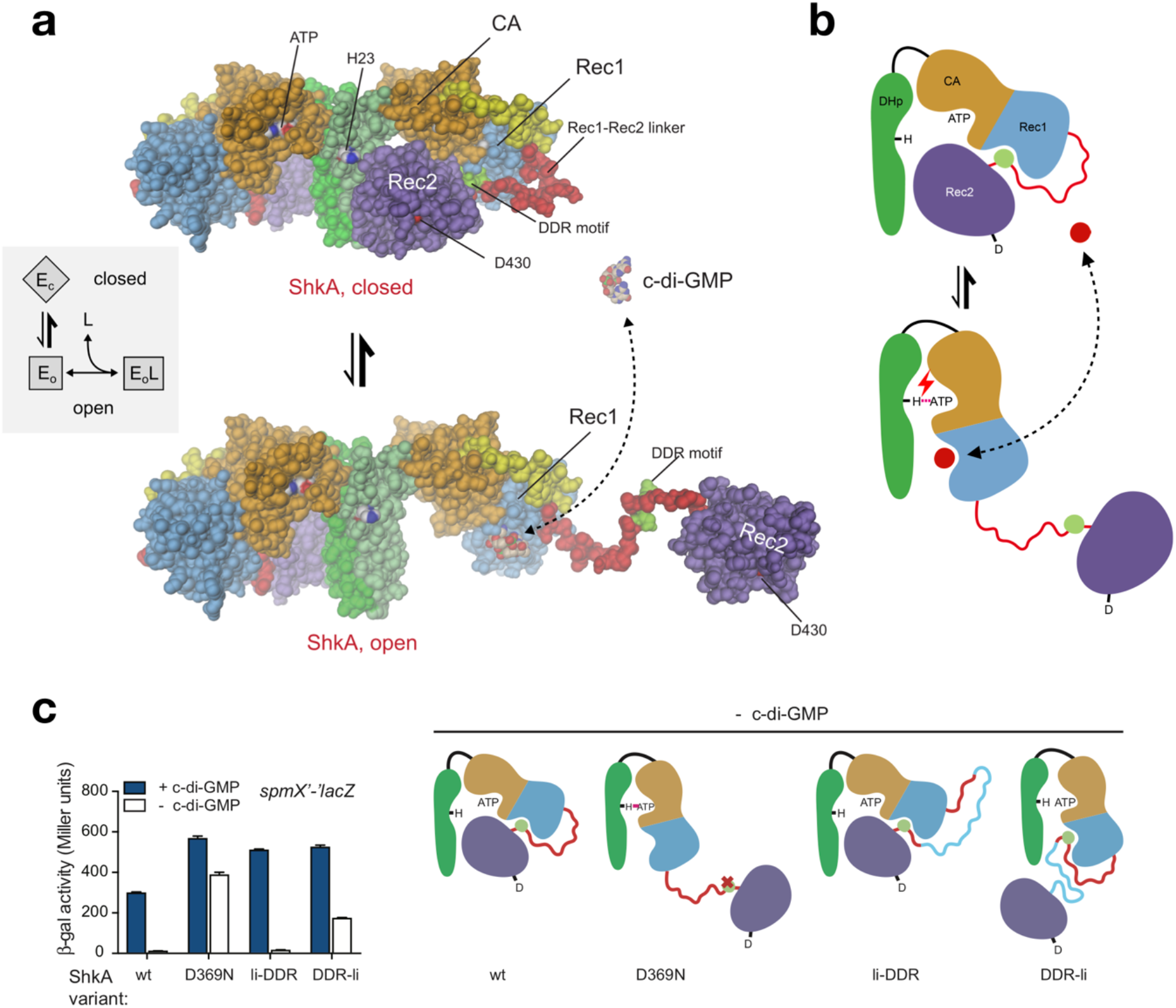
Mechanistic model of ShkA activation by c-di-GMP induced change of conformational equilibrium. **(a)** ShkA is in a dynamic equilibrium between a closed, auto-inhibited conformation as seen in the full-length crystal structure (top) and an ensemble of open, catalytically active conformations with liberated Rec2 domains. An arbitrarily modeled open conformation is shown at the bottom, full activity probably requires both subunits to be open. C-di-GMP can bind only to the open conformation, since, in the closed conformation, the binding site on Rec1 is obstructed by the DDR motif (light-green). Inset: Corresponding thermodynamic “ligand-coupled conformational equilibrium” (LCCE) model. **(b)** Schematic representation of the mechanism shown in (a). C-di-GMP (red circle) competes with the DDR motif (light-green circle) for Rec1 binding resulting in Rec2 liberation allowing catalysis (e.g. auto-phosphorylation as indicated). **(c)** *In vivo* activity of ShkA variants assayed in a strain harboring a *spmX*’-‘*lacZ* reporter. Mutation D369N affects a crucial Rec1 - Rec2 linker tethering interaction, mutants li-DDR and DDR-li have an elongated Rec1 - Rec2 linker with the insertion placed N- or C-terminal to the DDR motif as indicated in the scheme (right). Strains rcdG^0^ *ΔshkA* (- c-di-GMP) and *ΔshkA* (+ c-di-GMP) expressed the indicated *shkA* alleles from plasmid pQF. Note that no inducer (cumate) was present since leaky expression from the P_Q5_ promoter was sufficient to complement the *ΔshkA* phenotype. Mean values and standard deviations are shown (N=3).

The regulatory model was tested *in vivo* by analyzing the activity of ShkA variants with a β-Gal transcription assay. While in absence of c-di-GMP (rcdG^0^ strain), wt ShkA showed no transcriptional activity, introducing D369N or placing an insertion between the DDR motif and Rec2 (but not between Rec1 and DDR) rendered ShkA active (Fig. 4c). Thus, the results fully corroborate the regulatory model, as illustrated in the scheme (Fig. 4c, right). Additional support for the model is provided by AUC sedimentation velocity experiments (Supplemental Fig. 5) that show a decrease in the sedimentation coefficient upon ligand addition, indicative of a more open structure. In the following, the thermodynamic parameters of the LCCE model (inset to Fig. 4a) are quantified by functional investigations.

### ShkA auto-phosphorylation is c-di-GMP dependent, reversible and non-cooperative

To determine the reaction kinetics of the full-length ShkA and mutants thereof, we acquired progress curves of enzyme net phosphorylation (E*) by auto-radiography and of ATP to ADP turn-over by online ion exchange chromatography (oIEC, see Methods) quantification. First, we determined the kinetics of auto-phosphorylation separately using phosphotransfer deficient variants that had the phospho-acceptor D430 mutated (ShkA_D/A_) or the Rec2 domain deleted (ShkA_ΔRec2_). As found previously ^9,14^ and consistent with the proposed regulatory model, ShkA_ΔRec2_ is constitutively active (Figs. 5a, b), since it lacks the obstructing Rec2 domain. ShkA_ΔRec2_ attains very quickly (in less than 15 s) a stable phosphorylation state with or without the ligand. For ShkA_D/A_, the same fast kinetics is observed, but only in presence of c-di-GMP (Figs. 5c, d), which demonstrates that under this condition the mutant is fully active. In absence of the ligand, the reaction velocity is strongly reduced, but phosphorylation is obtained to almost the same degree as for the activated enzyme. The observations are consistent with the regulatory LCCE model (Fig. 4) under the assumption that the kinetics of the conformational equilibration are fast compared to the chemical reaction. Since the initial reaction velocity is at least 10-times larger in the presence of c-di-GMP (Fig. 5d), a lower boundary of 10 can be estimated for the conformational equilibrium constant K:= [E_c_]/[E_o_].

**Figure 5.**
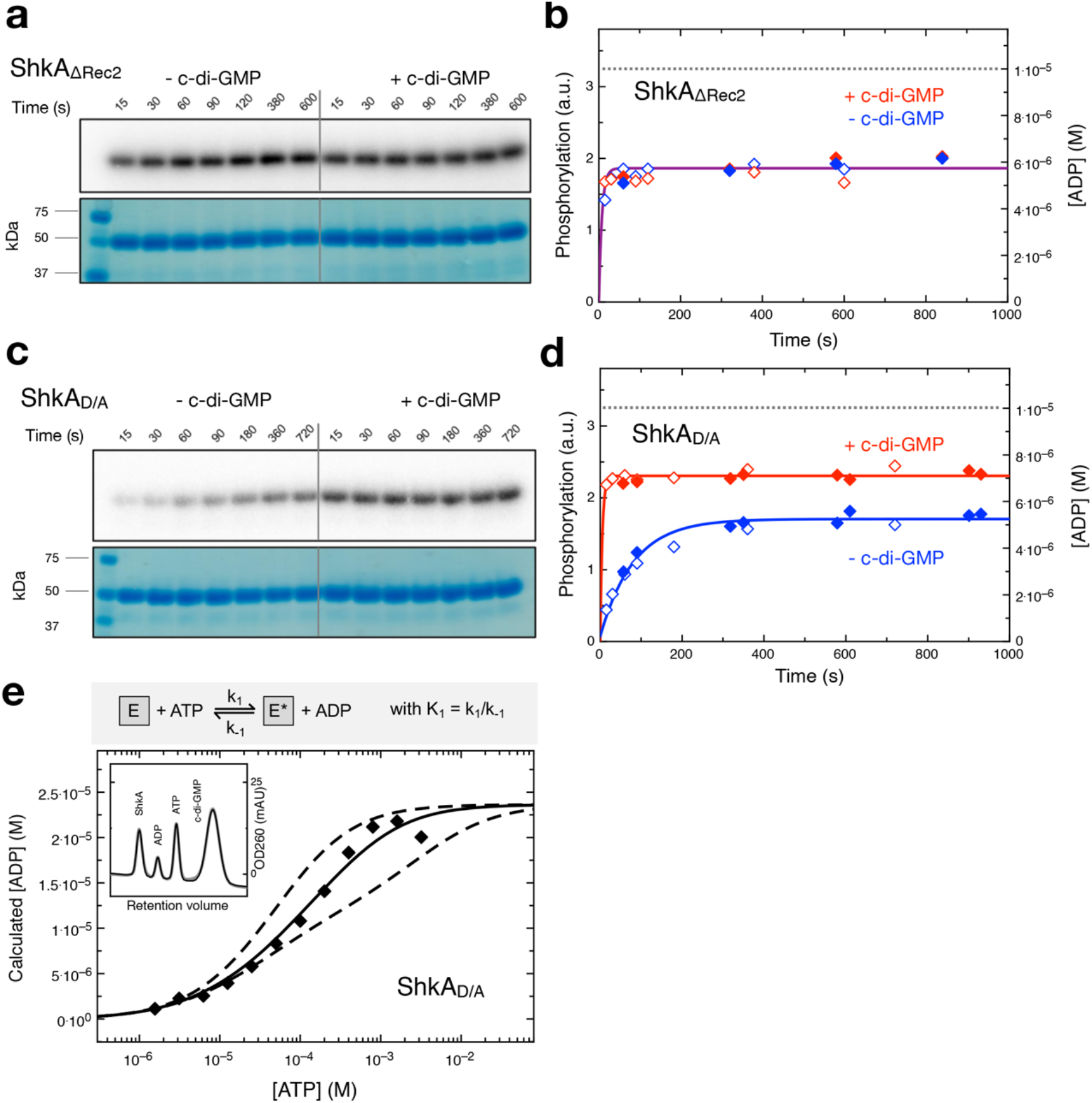
ShkA auto-phosphorylation is reversible and, in the presence of Rec2, controlled by c-di-GMP. **(a, c)** ShkA_ΔRec2_ and ShkA_D/A_ phosphorylation after indicated times of incubation as measured by autoradiography using [γ-^32^P]-ATP, in the absence and presence of c-di-GMP as indicated. SDS-PAGE gels are shown below and indicate equal loading. **(b, d)** Time course of ShkA_ΔRec2_ and ShkA_D/A_ phosphorylation (hollow symbols) and ADP production (filled symbols), in the absence (blue) and presence (red) of c-di-GMP. Phosphorylation values were calculated from the band intensities in panels (a, c), ADP concentrations were determined by oIEC. Solid lines represent exponential fittings, the dashed line indicates the employed enzyme concentration (10 μM). **(a-d)** [c-di-GMP]: 25 μM, [ATP]: 200 μM. **(e)** Equilibrium concentration of ADP as a function of initial ATP concentration as determined by oIEC. [ShkA_D/A_]: 20 μM, [c-di-GMP]: 50 μM. The continuous line is the fit of the simple reversible, non-cooperative bi-bi reaction shown on the top (see **Supplementary Fig. 6c**) to the data with K1 = k1/k-1 = 0.13. Reactions with positive or negative cooperativity would yield the dashed curves (calculated with equilibrium constants for second phosphorylation event of 1.3 and 0.013, respectively) that do not fit the data. Inset: Representative oIEC chromatogram.

To obtain more quantitative information about the auto-phosphorylation reaction, we measured ATP to ADP turnover by oIEC. Figures 5b and d show the ADP progress curves (filled symbols) of the phosphotransfer deficient ShkA mutants that turn out to be closely congruent with the appropriately scaled phosphorylation curves (open symbols). This indicates stable histidine phosphorylation at the employed condition as verified by long-term measurements (Supplemental Fig. 6b). Intriguingly, the progress curves (Figs. 5b, d) indicate that the phosphotransfer-deficient mutants do not proceed to full quantitative modification, i.e. did not reach the level corresponding to the enzyme concentration (10 μM). We reasoned that this may be due to reversibility of the auto-phosphorylation reaction, and that only a high ATP substrate concentration would shift the equilibrium completely to the product state. Indeed, upon ATP titration, the equilibrium ADP concentration was increased and reached the expected value close to the enzyme concentration indicating complete phosphorylation (Fig. 5e). A simple reversible bi-bi reaction model (see Methods) reproduced the data well and yielded an equilibrium constant K1 = k1/k-1 of 0.13 for activated ShkA_D/A_. Enzyme titration (Supplemental Fig. 6c) confirmed the result.

### C-di-GMP shifts thermodynamic ShkA equilibrium to the active state

Full-length ShkA is an ATPase, converting ATP to ADP *via* phospho-enzyme intermediates (Fig. 6a). Progress curves acquired at various ATP and ADP concentrations showed good agreement with a competitive product inhibition Michaelis-Menten model (Supplemental Fig. 7a, c). The derived Km and Ki parameters are both about 60 μM. These values agree reasonably well (within a factor of 2 to 3) with the ITC measurements (Supplemental Fig. 4). The derived effective rate constant k_eff_ = 0.21 s^−1^ is similar to the estimate of the auto-phosphorylation rate (0.43 s^−1^) that can be calculated from the ShkA_D/A_ data acquired in absence of c-di-GMP (Fig. 4d) assuming a K=49 (see below).

**Figure 6.**
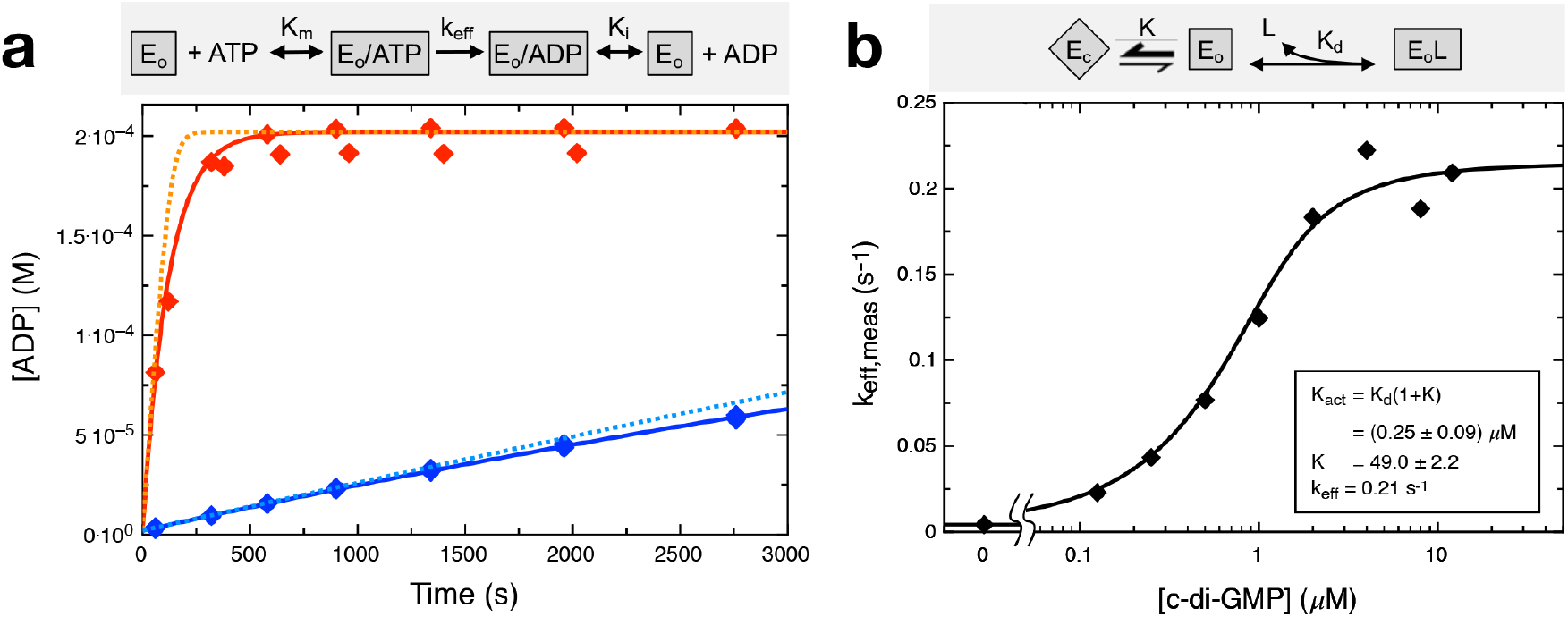
C-di-GMP activation profile of ShkA is consistent with a ligand-coupled conformational equilibrium (LCCE) model. **(a)** ADP production in the presence (red symbols) and absence of 25 μM c-di-GMP (blue) as measured by oIEC at room temperature. The reaction was started by addition of 200 μM ATP to 10 μM ShkA in reaction buffer. The data were fitted (solid line) with a competitive product inhibition Michaelis-Menten model (gray box; shown fully in **Supplementary Fig. 7a**) with KM and KI determined by ATP and ADP titration, see **Supplementary Figs. 7b-c**. With KI set to zero, the fit to the data is worse (dotted line) indicating significant product inhibition. **(b)** C-di-GMP activation profile obtained by acquisition of ADP progress curves at various c-di-GMP concentrations (**Supplementary Fig. 7b**) to yield keff,meas. The data were fitted to the LCCE model shown in the gray box, see Methods, Figure 7. Proposed domain arrangements during catalytic cycle of ShkA.

To quantitatively describe the activating effect of c-di-GMP, progress curves were acquired at various c-di-GMP concentrations (Supplemental Fig. 7b) and fitted with the Michaelis-Menten model. The resulting k_eff,meas_ values yielded the sigmoidal activation profile shown in Fig. 6b. The LCCE model fits the profile well and yields a conformational equilibrium constant K = 49 and an activation constant K_act_ = 0.29 μM, which agrees well with the (apparent) dissociation constant of c-di-GMP as determined by ITC in the presence of AMPPNP (1.5 μM, Fig. S5). The microscopic dissociation constant in the absence of the competing DDR motif can be calculated to K_d_ = K_act_/(1+K) = 6 nM. This value is smaller than those obtained for the DHp-CA-Rec1 construct by fluorescence competition and ITC (80 nM, 300 nM; Fig. 3e), which may be due to residual binding competition by the truncated Rec1-Rec2 linker.

In summary, the enzymatic characterization shows that, in the presence of c-di-GMP, ShkA efficiently catalyses auto-phosphorylation and subsequent phospho-transfer and dephosphorylation. The data fully support the regulatory LCCE model and thus confirm that the c-di-GMP activation profile is governed not only by the affinity of the ligand to its binding site, but also by the free energy difference between the conformational states.

### Structural modelling suggests a two-state catalytic cycle for hybrid histidine kinases

In a next step, we modelled the full catalytic cycle of ShkA based on the available structural and functional data and on general considerations of HHKs. Evidence has accumulated that the autophosphorylation competent form of HKs is an asymmetric dimer^6^. Recently, it has been proposed that such an asymmetric dimer can in fact catalyze both auto-phosphorylation and phosphoryl-transfer in concert and a structural model has been put forward for CpxA^17^. The corresponding ShkA model of an asymmetric dimer relieved from auto-inhibition is shown in Fig. 7a (top) where domains have been placed on the basis of homologous structures with competent auto-phosphorylation or phosphotransfer domain arrangement. Canonical auto-phosphorylation and phospho-transfer mechanisms can be anticipated, since the functional interfaces show no severe clashes or polarity mismatch.

**Figure 7.**
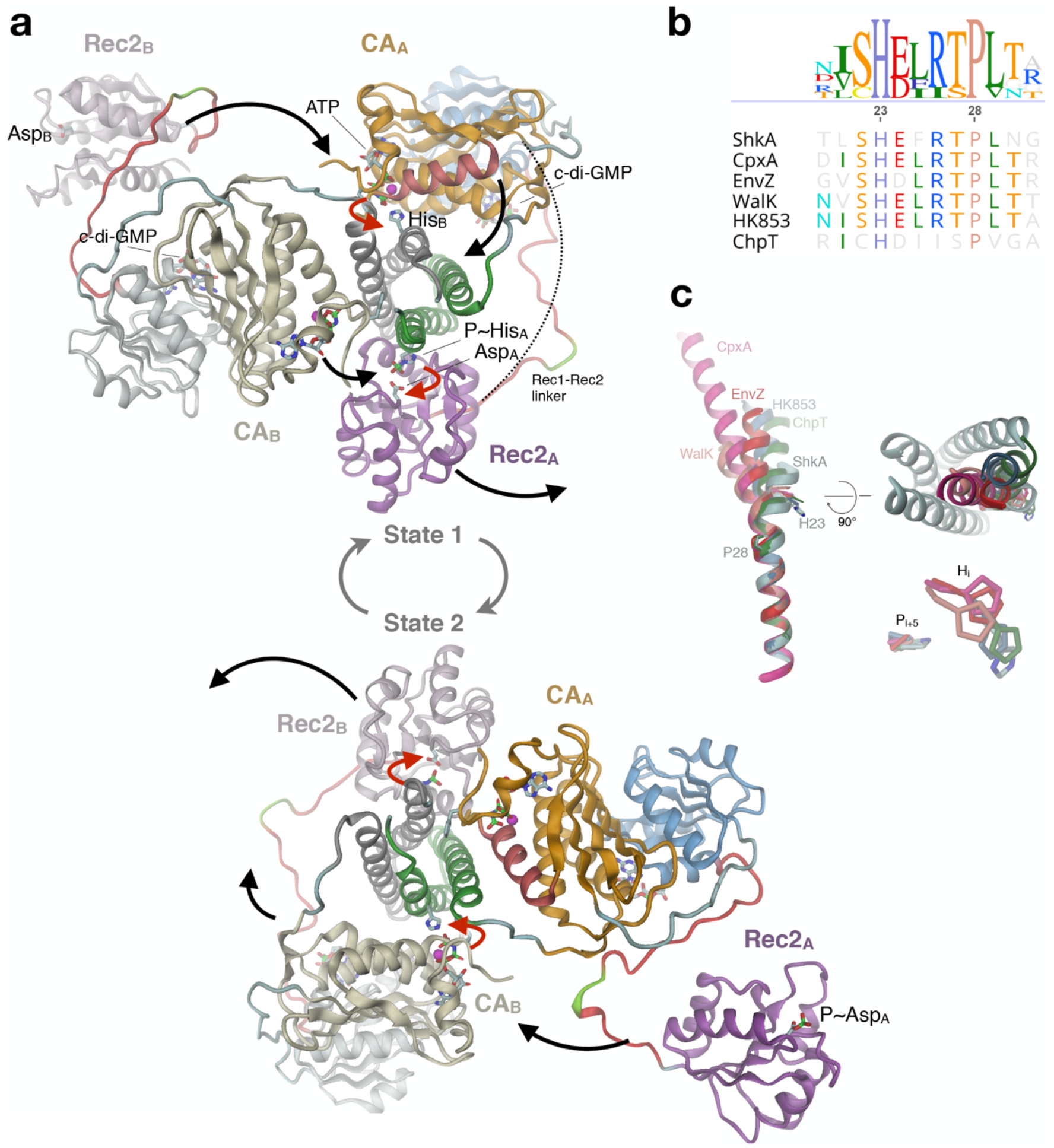
Proposed domain arrangements during the catalytic cycle of ShkA. **(a)** During the catalytic cycle, the enzyme adopts two asymmetric dimer states that are structurally identical, but with domain roles interchanged. Subunits are distinguished by color hue (A: strong, B: weak), the active His and Asp and ligands are shown in full. **State 1 (top)**: CA/ATP and Rec2 of chain A are poised for trans-autophosphorylation and cis-phosphotransfer, respectively. After concerted execution of the reactions (red arrows), CA/ADP and P~Rec2 dislodge and CA/ATP, Rec2 of chain B engage with the DHp histidines (black arrows) to yield **state 2 (bottom)**. The competent DHp/CA and DHp/Rec2 domain arrangements have been modelled based on the structures of CpxA (4biw^26^) and ChpT/CtrA(Rec) (4qpj^27^), respectively. In canonical HHK, CA would be linked directly to Rec2 (dashed line). **(b)** Part of DHp α1 sequence alignment of selected pfam HisKA family members. **(c)** Overlay of DHp α1 helix structures of the HKs of panel b after superposition of C-terminal parts. The structures with reported auto-phosphorylation competent domain arrangement (CpxA, 4biw; WalK, 5c93; EnvZ, 4kp4) and those in phosphotransfer or dephosphorylation/arrangement (HK853, 3dge; ChpT, 4qpj) and ShkA form structural clusters distinguished by the degree of helix kinking. Right: The kinase competent structures have the N-terminal part of α1 bent towards the center of the DHp bundle (ShkA in grey).

In the modelled asymmetric dimer, both CA and Rec2 of subunit A (CAA, Rec2A; bright colors) would be engaged in concerted catalysis resulting in a net transfer of a phosphoryl group from ATP to the active aspartate of Rec2A (red arrows), while the corresponding domain of subunit B (tinted colors) would be in stand-by mode. Subsequently, both domains would dislodge (black arrows), with CAA moving to a near-by parking position and Rec2A becoming available for down-stream phosphoryl transfer (Fig. 7a, bottom). Concurrently, the catalytic domains of subunit B would move to catalytically competent positions, resulting in the same structure as at the beginning, though 2-fold related and with the roles of the subunits interchanged (state 2). After concerted catalysis, equivalent domain rearrangements would bring back the enzyme to state 1.

In line with the left-handed connectivity of the two DHp helices, the model predicts that autophosphorylation occurs in *trans*^18^. Due to the almost equal distance between either of the C-termini of the Rec1 domains and the N-terminus of the competently docked Rec2 domain (about 60 Å), phosphotransfer may occur in *trans* or in *cis* as modelled. In any case, the Rec1 - Rec2 linker of ShkA is long enough (27 residues) to allow simultaneous execution of auto-phosphorylation and phosphotransfer. Notably, also for canonical HHKs (without an intervening domain like Rec1) this appears possible, since their CA - Rec linkers appear long enough (23 +/- 3 residues for the 23 HHKs in *C. crescentus*) to bridge the distance (again 60 Å, dashed line in Fig. 7a).

## Discussion

The structural, dynamic, and functional results presented in this study consistently show that activation of ShkA proceeds *via* the c-di-GMP mediated liberation of a locked, auto-inhibited state (Fig. 4). Thereby, c-di-GMP competes with the tethering of a domain linker (Figs. 3a,d) to unleash the C-terminal domain allowing the enzyme to step through the catalytic cycle that involves large domain motions (Fig. 7). C-di-GMP interference with protein - protein interactions as a regulatory mechanism was also proposed for a YajQ-like transcription factor / co-activator complex^19^, but structural information is missing.

In ShkA, c-di-GMP binds to the edge strand (β5) of the Rec1 β-sheet and the following β5 - α5 loop. The Watson-Crick edges of both guanines form H-bonds with the main-chain (Fig. 3b), reminiscent of the backbone interactions in a β-sheet. The same kind of interaction occurs in the CckA/c-di-GMP complex, where one of the guanines binds to the β-sheet edge of the CA domain^13^. In the accompanying study^9^, Y338 was identified as a c-di-GMP binding residue in a targeted alanine scan. In addition to Y338A, several other mutants (R324A, I340A, S347A, and Q351A) showed severely impaired activity *in vivo*^9^. Since all these residues contribute to c-di-GMP binding (Fig. 3b), this can be attributed to an impaired activation by the cellular c-di-GMP pool and suggests that the affinity of wild-type ShkA to c-di-GMP is finely tuned to its raise in concentration during cell cycle progression^20^.

A mutation of R344, the remaining c-di-GMP binding residue, showed a wild type-like phenotype. This may be explained by the dual role of R344 being involved in c-di-GMP binding as well as Rec1 - Rec2 linker tethering (Figs. 3a, d), resulting in opposing effects on activation upon mutation. Although R324 also has such a dual role, its strong phenotype suggests that this residue contributes more to c-di-GMP binding than to linker tethering. These observations demonstrate that identification of ligand binding residues is not straightforward if they are also involved in opposing activities like protein - protein interactions. However, mutation of the DDR motif or inserting a loop between the motif and the Rec2 domain clearly resulted in constitutively active enzyme and, thus, confirmed the importance of Rec2 tethering for auto-inhibition (Fig. 4c).

The NMR studies on full-length ShkA using specific isotope labelling of the isoleucine methyl groups provided atomic scale information of structural and dynamic changes upon ligand binding (Fig. 2). Binding of c-di-GMP, but not AMPPNP, changed the dynamics of Rec2 substantially and exclusively, leading to the conclusion that the mobility of the Rec2 domain is restricted in the absence of c-di-GMP but is drastically enhanced upon addition of the ligand such that its relaxation times become comparable to those of an individual protein of this size. The binding of c-di-GMP to Rec1 therefore releases Rec2 allosterically.

In order to relate the mechanistic “activation by domain liberation” model of ShkA with the function of this hybrid kinase, comprehensive enzymatic data were acquired. The newly developed oIEC method proved highly valuable as it enabled the efficient recording of quantitative enzyme progress curves. We found, using phospho-transfer deficient mutants, that ShkA auto-phosphorylation is clearly reversible. Under various titration regimes, the equilibrium concentrations were consistent with a simple reversible bi-bi reaction with the equilibrium largely on the side of the reactants (Fig. 5e, Supplemental Fig. 6c). Thus, the degree of phosphorylation depends on the mono-nucleotide concentrations, and a high ADP concentration promotes enzyme dephosphorylation by back transfer of the phosphoryl group onto ADP. Reversibility of the HK auto-phosphorylation reaction, first demonstrated by Jiang et al. in their pioneering study on NRII^21^, is probably a general feature of ATP driven histidine phosphorylation, but has been largely ignored in the literature. It should be tested whether it can explain the ADP dependence of the kinase reaction as observed for HK853^22^ and other HKs^23^ and whether this feature is of physiological relevance. The rate constant of autophosphorylation is with k_1_ = 0.4 s^−1^ at least 20 times faster than measured for other HKs^13,24^. Possibly, some of these enzymes were partly impaired by truncation and/or not fully activated. ATP turnover of full-length ShkA proceeds somewhat slower, which demonstrates that phospho-transfer and P~Asp hydrolysis are not much slower. To answer whether the latter reaction is entirely due to the intrinsic lability of the phospho-aspartate or catalysed by docking of the Rec2 domain to the DHp bundle^3,15^ requires further investigations.

It is generally believed that only one of the two symmetry related histidines of HKs can get phosphorylated at a given time, since the kinase state involves an asymmetric dimer. Most likely the asymmetry is rooted in the inward kinking of the N-terminal part of DHp α1 helix upon interaction with the ATP complexed CA domain (Fig. 7c), which would be sterically incompatible with an analogous motion of α1 of the other subunit. This has been discussed and analyzed thoroughly by Bhate and coworkers^6^ and corroborated by the recent structure and modelled dynamics of WalK^25^. Such asymmetric catalysis is not necessarily equivalent with half-of-sites modification and is, therefore, not at odds with our finding of complete modification at high ATP concentration (Fig. 5e). Disengagement of the CA domain after the first phosphorylation event could easily allow the other CA domain to approach the second histidine for modification (compare states 1 and 2 in Fig. 7a). Intriguingly, however, CpxA shows a hemi-phosphorylated crystal structure in the presence of ATP^17^, although the non-phosphorylated histidine seems perfectly poised to react with CA bound ATP in the observed canonical kinase competent arrangement. Based on our thermodynamic results on ShkA, we predict that CpxA behaves similarly, with the equilibrium of the reversible auto-phosphorylation reaction being on the side of the reactants (ATP, His). This would explain why the substrate and not the product complex was observed, since the products (ADP, P~His) would react back and ADP would stay in the active site assuming strong crystal lattice constraints.

Our analyses provide mechanistic and kinetic insight into how c-di-GMP competes with a proteinprotein interaction to activate an HHK, and how ADP affects the auto-phosphorylation reaction. For a full mechanistic understanding of HHKs, more investigations are required to decipher the kinetics of the phospho-transfer and auto-dephosphorylation steps and to unravel the physiological advantage of combining histidine kinase core and cognate receiver domain on one polypeptide.

## Accession codes

Atomic coordinates and structure factors for the ShkA/AMPPNP and the ShkA_Rec1_/c-di-GMP complexes have been deposited in the Protein Data Bank under accession numbers 6QRJ and 6QRL, respectively. Sequence-specific backbone resonance assignment of the ShkA_Rec2_ domain has been submitted to the Biological Magnetic Resonance Data Bank under accession code 27882.

## Acknowledgments

We thank the beamline staff at the Swiss Light Source in Villigen for expert help in data acquisition and T. Sharpe from the Biophysics facility at the Biozentrum Basel for expert biophysical support. We thank A. Eberhardt and A. Roulier for technical assistance, and H. Pickersgill (Life Science Editors) for editorial assistance.

## Author Contributions

B.N.D., E.A., R.B., T.S., I.P-M., S.H., U.J. designed the experiments. B.N.D. and E.A. purified and crystallized ShkA and ShkARec1. B.N.D. solved and analyzed the crystal structures and generated the models describing the mechanisms. R.B. and S.H. performed and analyzed the NMR experiments. E.A., B.N.D., F.M., I.P-M., A.K. and C.v.A. performed biochemical and biophysical experiments. E.A., B.N.D., T.S. analyzed the biophysical and biochemical data and performed the bioinformatic analyses. T.S. and E.A. implemented the kinetic models. A.K. designed and performed the transcription assays. T.S., B.N.D., E.A., R.B., I.P-M., S.H., U.J. wrote the manuscript with contributions from all other authors. Funding acquisition, T.S., U.J., S.H.

## Methods

### Cloning and protein expression for *in vitro* experiments

Vectors, primers, and recombinant proteins used in this study are listed in **Supplementary Tables 1, 2, and 3**. *In vitro* experiments presented in this study were carried out using protein expressed from a pET-28a-shkA plasmid. The pET-28a-shkA vector carrying the gene coding for full-length, wild-type ShkA (Uniprot: Q9ABT2) was the same as used in the preceding study^9^. Most point mutants and wild-type variants were generated from this vector following the Q5 Site-Directed Mutagenesis Kit protocol (E0554, New England Biolabs) and primers 1-8 (Sup. Table 2). The results were verified by sequencing. Isoleucine point mutants used for the NMR study were cloned by standard mutagenesis using Phusion polymerase (Thermo Fisher Scientific) and primers 9-16 (Sup. Table 2), and the results verified by sequencing. The *E. coli* DH5α strain (Invitrogen) was used for cloning purposes. Bacteria were allowed to grow in lysogeny broth (LB; 10 g tryptone, 5 g yeast extract, 10 g NaCl) supplemented with 50 μg/ml Kanamycin (LB-Kan) and LB-Kan-containing agar plates at 37 °C. After successful cloning, pET-28a-shkA plasmids were extracted from *E. coli* DH5α cells following QIAprep Spin Miniprep Kit protocol (QIAGEN) and used to transform the expression strains *E. coli* BL21(DE3) or similarly efficient Rosetta cells (Novagen). For protein expression, adequate amounts of LB-Kan media were inoculated with 1% pre-culture of transformed cells. Grown cultures were induced with 1 mM isopropyl 1-thio-β-D-galactopyranoside (IPTG) at an OD600 of 0.6 – 0.8. The incubation temperature was reduced to 22 °C for overnight protein expression. Cells were harvested by centrifugation at 9’000 RCF for 10 minutes at 4 °C. Pellets were stored at −80 °C or lysed immediately.

### Protein purification

Purification was entirely performed at 4 °C. Pellets were homogenized in lysis buffer containing immobilized metal affinity chromatography (IMAC) loading buffer (500 mM NaCl, 30 mM Tris-HCl, 5 mM MgCl_2_, 20 mM Imidazole, pH 7.5), cOmplete EDTA-free protease inhibitors (Roche) and bovine pancreas DNase I (Roche). Cell lysis was performed either with the ultrasonicator (35 % amplitude, 7×30 s pulses with short interval) or the microfluidizer (two passes, 10’000 psi). The lysate was ultracentrifuged at 30,000 RCF for 30 min, to remove cell debris and suspended particles. The clear supernatant was applied on a 5 mL Ni-NTA column (GE Healthcare) pre-equilibrated with IMAC loading buffer. Bound protein was eluted with a linear gradient of IMAC elution buffer (500 mM NaCl, 30 mM Tris-HCl, 5 mM MgCl_2_, 0.5 or 1.0 M Imidazole, pH 7.5) using an ÄKTA Purifier system (GE Healthcare). Fractions containing the desired protein were pooled and concentrated to a volume of 5 mL or below. Depending on size, the sample was loaded on a HiLoad 26/60 Superdex 200pg or a Superdex 75pg gel filtration column pre-equilibrated with SEC buffer (100 mM NaCl, 30 mM Tris-HCl, 5 mM MgCl_2_, pH 7.5). The concentrations of the collected samples were quantified by UV absorption with a NanoDrop 2000 spectrophotometer (Thermo Fisher Scientific) and either used freshly (e.g. for crystallization) or stored at −80 °C.

### ShkA crystallization

Protein solubilised in SEC buffer (see above) was crystallised using the sitting-drop vapor diffusion method. Sets of 3-drop MRC plates were prepared with a Gryphon robot (Art Robbins Instruments). Full-length ShkA was crystallised at a concentration of 10 mg/ml in presence of AMPPNP and c-di-GMP (1:5:3 molar ratio) with a crystallization mixture consisting of 0.2 M DL-Glutamic acid monohydrate, 0.2 M DL-Alanine, 0.2 M Glycine, 0.2 M DL-Lysine monohydrochloride, 0.2 M DL-Serine, 0.1 M Sodium HEPES pH 7.5, MOPS (acid), 40 % v/v ethylene glycol and 20 % w/v PEG 8000 (Morpheus-H7, Molecular Dimensions) at room temperature. Surprisingly, the resulting structure did not contain c-di-GMP. Probably, the major fraction of molecules that had c-di-GMP bound and, thus, an open, mobile domain arrangement (see main text) did not crystallise, but rather the minor fraction of uncomplexed (c-di-GMP) molecules with their compact, locked domain arrangement. Crystals of ShkARec1 were obtained in the presence of c-di-GMP (1:3 molar ratio) at a concentration of 20 mg/ml in 0.2 M Ammonium sulfate, 0.1 M Bis-Tris pH 6.5 and 25% w/v PEG 3350 (SG01-F5, Molecular Dimensions) at 4 °C in cold room.

### Data collection and structure determination

X-ray diffraction data were collected at the Swiss Light Source (SLS), Villigen, Switzerland, at 100 K. Data were indexed, integrated, scaled, and merged using XDS^28^ and the CCP4i2 suite^29^. The crystal structure of full-length ShkA was solved by molecular replacement^30^, using a homology model of ShkA (CA) based on the CA structure of DivL (PDB code 4q20) as a first search model. After having placed the CA domain successfully, subsequent molecular replacement searches were conducted using a Chainsaw/CCP4^29,31^ model of Rec1 derived from the first Rec domain of 3luf with helix 3 removed. The Rec2 domain was localised with help of a Chainsaw model derived from the second receiver domain of the same PDB entry. At this time point, clear density was obtained for the DHp bundle and its model was obtained by molecular replacement with the individual α1 and α2 DHp helices of DivL (4q20). The ShkARec1 crystal structure was also solved by molecular replacement^32^ using the corresponding domain of the full-length ShkA structure. For both crystal structures, phases and models were further improved by automated model building using PHENIX AutoBuild^33^. Extensive additional manual model building was carried for the Rec1 - Rec2 linker containing DDR motif and the loop (connector) between DHp α1 and α2 in Coot^34^. Iterative rounds of model building and refinement in Refmac5^29^ resulted in a map of sufficient quality to place the missing parts of the peptide chain along with the ligands. Figures were prepared with Dino (http://dino3d.org).

### Nuclear Magnetic Resonance

All NMR spectra were recorded at 20 °C on a Bruker Avance-700 MHz spectrometer equipped with a cryogenically cooled triple-resonance probe. The following experiments were recorded with the ShkARec1 construct: 2D [^15^N,^1^H]-TROSY^35^, 2D [^13^C,^1^H]-HMQC, 3D HNCA, 3D HNCACB, 3D ^15^N-HSQC-^1^H,^1^H-NOESY, 3D ^13^C-HSQC-^1^H,^1^H-NOESY, 2D [^13^C,^1^H] HMQC ^13^C T_2_^36^; with the ShkA_Rec2_ construct: 2D [^15^N,^1^H]-TROSY, 2D [^13^C,^1^H]-HMQC, 3D HNCA, 3D HNCACB, 3D CBCA(CO)NH, 2D [^13^C,^1^H] HMQC ^13^C T_2_; with the full-length ShkA construct: 2D [^15^N,^1^H]- TROSY, 2D [^13^C,^1^H]-HMQC, 2D [^13^C,^1^H] HMQC ^13^C T_2_. Sequence-specific backbone assignment of ShkA_Rec2_ was obtained from standard triple resonance NMR experiments recorded in NMR buffer (30 mM Tris-buffer at pH 7.2 with 160 mM NaCl, 5 mM MgCl_2_ and 1 mM DTT). The sequence-specific backbone assignment of ShkARec1 is reported in Kaczmarczyk et al.^9^ (BMRB 27768). Assignment of the δ1 isoleucine methyl groups of ShkARec1 was obtained from 3D ^15^N-HSQC-^1^H,^1^H-NOESY and 3D ^13^C-HSQC-^1^H,^1^H-NOESY NMR experiments by identifying unambiguous NOEs consistent with the crystal-structure of ShkA bound to AMPPNP (PDB:6QRJ). Methyl group assignments were transferred from the isolated ShkARec1 domain to full-length ShkA by spectral comparison. Additional isoleucine δ1 methyl group assignments of ShkA were obtained using mutagenesis (I259V, I340V, I390V, I405V).

NMR experiments of isolated ShkARec1, ShkA_Rec2_ domains and ShkA full-length protein and its ligand complexes with AMPPNP and/or c-di-GMP were recorded in NMR buffer at 20 °C. 2D [^15^N,^1^H]-TROSY and 2D [^13^C,^1^H]-HMQC spectra were recorded with [U-^2^H,^15^N]-Ile δ1-[^13^C,^1^H]-labelled ShkA protein of the following concentrations: 270 μM apo ShkA; 270 μM ShkA + 1.2 mM AMPPNP; 270 μM ShkA + 1.2 mM c-di-GMP and 270 μM ShkA + 1.2 mM AMPPNP + 1.2 mM c-di-GMP.

^13^C T_2_ transverse relaxation times of isoleucine δ1 methyl groups were measured using [Ile δ1- ^13^C,^1^H]-labelled sample of ShkARec1, ShkARec1 bound to c-di-GMP, ShkA_Rec2_, full-length ShkA bound to AMPPNP and full-length ShkA bound to AMPPNP and c-di-GMP using the 2D [^13^C,^1^H] HMQC ^13^C T_2_ experiment. The experimental conditions were the same as for the binding experiments. Spectra were recorded with different T delays (Rec1: 3.1, 46.8, 93.6, 140.4, 187.2, 374.4, 468 ms; Rec2: 3.1, 31.2, 62.4, 93.6, 124.8, 156, 187.2, 312, 468 ms; full-length ShkA (1): 3.1, 18.7, 37.4, 74.9, 149.8 ms; full-length ShkA (2): 3.12, 15.6, 31.2, 46.8, 62.4, 124.8, 187.2 ms) and the peak intensities were fitted to an equation of the form: Peak Intensity = A exp(-T/T_2_). The error for T_2_ was derived from the fit of the data by bootstrapping for the isolated domains and as the standard deviation for two independent measurements for full-length ShkA.

### Isothermal Titration Calorimetry (ITC)

Measurements were performed with a MicroCal VP-ITC calorimeter (Malvern Instruments) at 25 °C. Samples were diluted in ITC buffer (100 mM NaCl, 20 mM Tris-HCl, 5 mM MgCl_2_, 1 mM DTT, pH 7.5). A total of 30 injections of 10 μl each (besides the first injection with 2 μl) were made with a spacing time of about 400 seconds. Thermograms were analysed with AFFINImeter^37^ (S4SD, Santiago de Compostela, Spain) using a 1:1 binding model with the stoichiometry fixed to 1. Tested ligands: AMP-PNP (Sigma Aldrich), ADP (Sigma Aldrich), c-di-GMP (Biolog, Bremen, Germany).

### Fluorometric binding competition assay

The dissociation constant of 2’-Fluo-AHC-c-di-GMP (fluo-c-di-GMP; Biolog, Bremen, Germany) to full-length ShkA and its variant as determined by monitoring the change in fluorescence intensity upon protein titration. This was followed by c-di-GMP titration to protein in presence of fluo-c-di-GMP to compete out the labelled ligand. Data were fitted to the competitive ligand binding model given by Z-H. Wang^38^ to obtain the dissociation constant of unlabelled c-di-GMP. Experiments were performed with BioTek Synergy H1 and 2 plate readers. Fluorescence intensity was recorded at either λ_em_ = 517 nm with λ_ex_ = 494 nm (Synergy H1), or λ_em_ = 516/20 nm with λ_ex_ = 485/20 nm (bandpass filters, Synergy 2), in both cases at a height of 8.5 mm and sensor gain of 80-100. Measurements were performed in ITC buffer.

### Sedimentation velocity analytical ultracentrifugation (SV-AUC)

Sedimentation velocity was measured with a Beckman Coulter XL-I ultracentrifuge with interference optics. Experiments were performed with 12 mm double-sector centerpieces, 8-hole rotors (Beckman An50Ti) at 42 krpm and 25 °C. Samples (1 mg/ml = 18 μM) were prepared in SEC buffer. Viscosity (0.901 cp) and density (1.0035 g/ml) of the buffer were measured separately at 25 °C with an Anton-Paar instrument. Analysis of the raw data was performed using SEDFIT^39^. The “Continuous c(S) distribution with bimodal f/f0” fitting model was used to mitigate the effect of small molecules/contaminants present in the samples.

### β-Gal assay

Plasmids for β-Gal assays were constructed as follows. A fragment encoding the N-terminal predicted unstructured part of RPA4224, a predicted ortholog of the anti-sigma factor NepR^40^, was PCR-amplified from *Rhodopseudomonas palustris* CG009 genomic DNA with primers 17/18 (pQF-shkA-liDDR; Sup. Table 2) or 19/20 (pQF-shkA-DDRli; Sup. Table 2), digested respectively with PstI or MluI (New England Biolabs) and cloned in pQF-shkA predigested with either restriction enzyme. The correct orientation of the inserted fragment was verified by sequencing. Plasmids pQF-shkA and pQF-shkA(D369N), and the parental plasmid pQF, were described previously^9,41^. Strains (AKS297 or UJ9691) harboring the pAK502-spmX reporter plasmid^9^ were transformed with pQF-shkA, pQF-shkA(D369N), pQF-shkA-liDDR or pQF-shkA-DDRli by electroporation. Single colonies were inoculated in 2 ml PYE supplemented with chloramphenicol (1 μg/ml) and oxytetracyline (1 μg/ml) and grown overnight at 30°C in a drum roller. Cultures were diluted the next day 20-fold in 2 ml of the same medium, followed by further incubation for an additional 4.5 h under the same conditions before sampling. β-Gal assays were essentially performed as described before according to the method of Miller^42^.

### Protein phosphorylation assay

Net-phosphorylation of full-length ShkA and its variants were analysed by auto-radiography as described elsewhere^9^. In short, reactions were run in Enzymatic Reaction buffer (30 mM Tris-HCl, pH 7.5, 50 mM NaCl, 50 mM KCl, 5 mM MgCl_2_) in the presence of 200 μM ATP (Sigma Aldrich) and 5 μCi [γ-32P]-ATP (3,000 Ci mmol^−1^, Hartmann Analytic) either at room temperature or at 4 °C. Additional nucleotides were added at indicated time points. Reactions were interrupted with SDS sample buffer (Lämmli) and subsequently loaded on precast 4-20 % gradient SDS PAGE gels (BIORAD). Wet gels were exposed to phosphor screen (0.5–3 hrs) before being scanned using a Typhoon FLA 7000 imaging system (GE Healthcare). Background subtracted autoradiograph band intensities were determined with ImageStudio (Li-COR, Lincoln, Nebraska USA).

### <u>O</u>nline <u>i</u>on <u>e</u>xchange <u>c</u>hromatography (oIEC) assay for acquisition of nucleotide turn-over progress curves

Nucleotide concentrations of an ongoing enzymatic reaction were determined in real-time by fast protein liquid chromatography (FPLC) using an ÄKTA Purifier instrument (GE Healthcare) equipped with a 1 mL Resource Q anion exchange (AEX) column and an A-905 autosampler.

With the newly developed oIEC method we acquired automatically quantitative enzyme progress curves by repetitive cycles of sample loading followed by salt gradient elution. Thereby, the reaction was started by addition of ATP substrate to the reaction mix in reaction buffer at t=0. This was followed by sequential aspiration of aliquots to the column in loading buffer (20 mM Tris-HCl, pH 8.0) at defined time points. Note that when the sample arrives on the column the reaction is stopped without the need of further intervention due to substrate immobilization. Elution was achieved by a linear gradient from 0 - 1 M (NH4)2SO4. The absorbance at 260 nm was constantly monitored using a UV-900 monitor (GE Healthcare). The chromatogram peaks corresponding to the nucleotides were fitted by Gaussians using a custom made automized routine implemented in ProFit 7 (QuantumSoft, Uetikon am See, Switzerland). Peak areas (mAU · ml) were converted to molar nucleotide amounts using a scale factor obtained by calibration with a set of serially diluted ATP samples.

### Thermodynamic modelling of reversible autophosphorylation

The equilibrium constant K of reversible protein (E_o_) auto-phosphorylation (inset to Fig. 4B) is given by

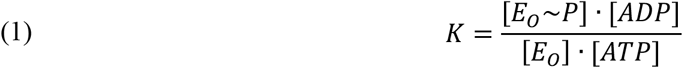

To solve the reaction for given total enzyme (E_tot_) and ATP (ATP_tot_) concentrations, we consider the following (non-equilibrium) starting concentrations

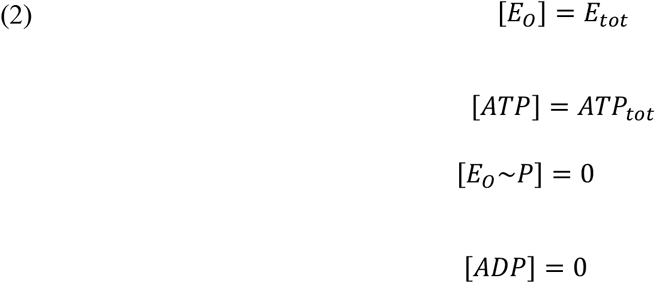

which will reach at equilibrium

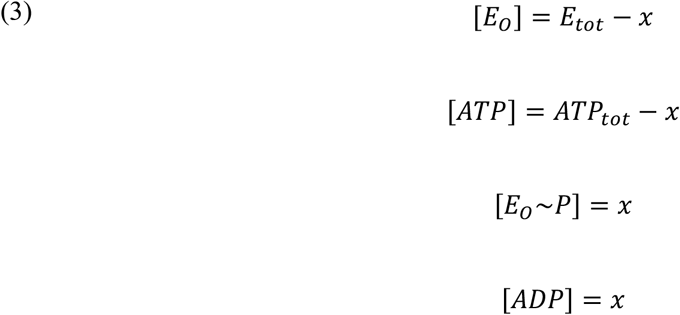

Inserting (3) into (1) yields

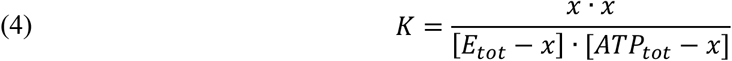

which can be solved analytically (WolframAlpha, http://www.wolframalpha.com) for x. Substituting x into equations 3 yields the four equilibrium concentrations.

In case of a dimeric enzyme with two phosphorylation sites, four reactions have to be considered

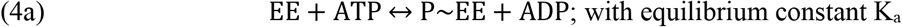

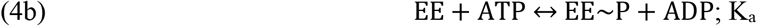

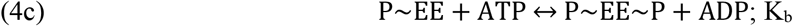

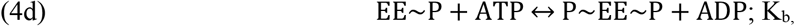

whereby attachment of the first phosphoryl group (reaction 4a or 4b) would be governed by equilibration constant K_a_, and attachment of the second phosphoryl group (reaction 4c or 4d) by Kb. Positive or negative cooperativity would be manifested by K_b_>K_a_ and K_b_<K_a_, respectively. No cooperativity would be equivalent to K_b_ = K_a_.

### Thermodynamic modelling, ligand induced change in equilibrium (LICE)

For an enzyme in dynamic equilibrium between two states (active E, inactive C) and a ligand L that binds exclusively to the E-state (inset to Fig. 5B), the following equations can be formulated

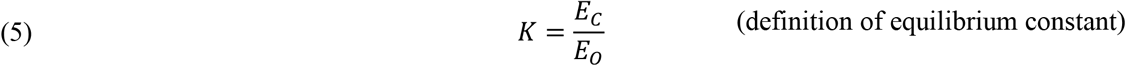

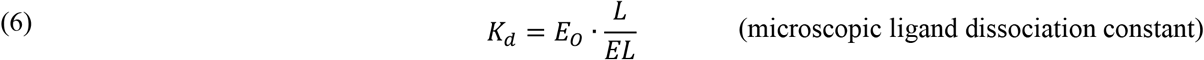

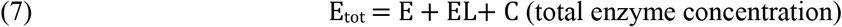

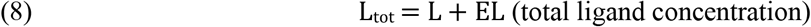

This set of four linear equations (5-8) can be solved analytically (WolframAlpha, http://www.wolframalpha.com) for the four unknown concentrations. The fraction of active enzyme molecules, assuming that ligand binding does not interfere with activity, is then given by:

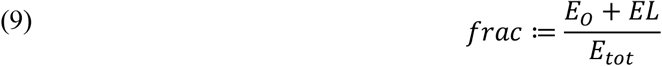

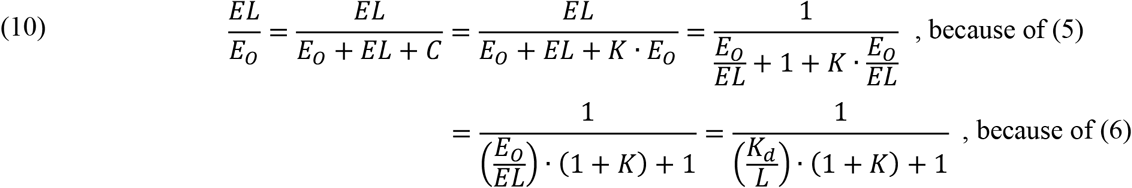

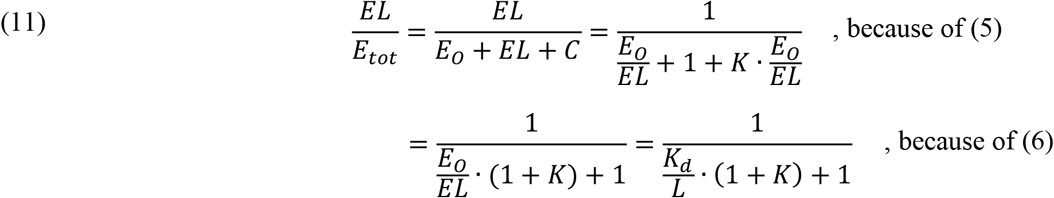

which is the form of a hyperbolic activation curve

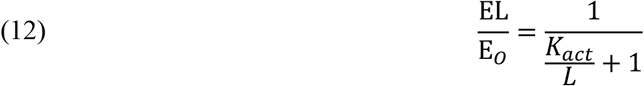

with an activation constant of

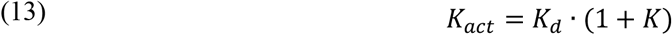

The measured (apparent) rate constant is then given by

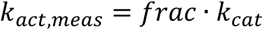

