## Supplementary Tables for "Hybrid histidine kinase activation by cyclic di-GMP-mediated domain liberation"

| ID | Name | Description | Construction | Reference |
|---|---|---|---|---|
| 1 | pET28a-shkA | Vector pET-28a(+) with shkA gene, KanR. To express N-terminal polyHis fusion of ShkA. | N/A | Kaczmarczyk et al., 2019 |
| 2 | pET28a-shkA-Rec1 | To express N-term. polyHis fusion of $ShkA_{Rec1}$. | N/A | Kaczmarczyk et al., 2019 |
| 3 | pET28a-shkA-Rec2 | To express N-term. polyHis fusion of $ShkA_{Rec2}$. | NEB Q5 SDM Kit, with primers 1 and 2 | This paper |
| 4 | pET28a-shkA-DHp-CA-Rec1-DDR | To express N-term. polyHis fusion of $ShkA_{DHp\text{-}CA\text{-}Rec1\text{-}DDR}$. | NEB Q5 SDM Kit, with primers 3 and 4 | This paper |
| 5 | pET28a-shkA-DHp-CA-Rec1 | To express N-term. polyHis fusion of $ShkA_{DHp\text{-}CA\text{-}Rec1}$, also known as $ShkA_{\Delta Rec2}$. | NEB Q5 SDM Kit, with primers 5 and 6 | This paper |
| 6 | pET28a-shkA(I259V) | To express N-term. polyHis fusion of $ShkA_{I259V}$. | with primers 9 and 10 | This paper |
| 7 | pET28a-shkA(I340V) | To express N-term. polyHis fusion of $ShkA_{I340V}$. | with primers 11 and 12 | This paper |
| 8 | pET28a-shkA(I390V) | To express N-term. polyHis fusion of $ShkA_{I390V}$. | with primers 13 and 14 | This paper |
| 9 | pET28a-shkA(I405V) | To express N-term. polyHis fusion of $ShkA_{I405V}$. | with primers 15 and 16 | This paper |
| 10 | pET28a-shkA(D430A) | To express N-term. polyHis fusion of $ShkA_{D430A}$. | NEB Q5 SDM Kit, with primers 7 and 8 | This paper |
| 11 | pQF-shkA | Vector pQF with shkA gene, to express ShkA. For *in vivo* β-Gal assay | N/A | Kaczmarczyk et al., 2019 |
| 12 | pQF-shkA(D369N) | To express $ShkA_{D369N}$, for *in vivo* β-Gal assay | N/A | Kaczmarczyk et al., 2019 |
| 13 | pQF-shkA-liDDR | To express $ShkA_{liDDR}$, for *in vivo* β-Gal assay | with primers 17 and 18 | This paper |
| 14 | pQF-shkA-DDRli | To express $ShkA_{DDRli}$, for *in vivo* β-Gal assay | with primers 19 and 20 | This paper |

**Supplemental Table 1 |** Plasmids used to express the proteins used in this study. Primer numbers refer to Suppl. Table 2.

| ID | Name | Sequence (5’>3’) | Length | Tm* (°C) | Exp Ta** (°C) |
|---|---|---|---|---|---|
| 1 | Q5SDM_ShkARec2_F | CCC GCC CAC GAC GAC CGC | 18 | 67 | 70 |
| 2 | Q5SDM_ShkARec2_R | ATG GCT GCC GCG CGG CAC | 18 | 65 | 70 |
| 3 | Q5SDM_ShkADHp-CA-Rec1-DDR_F | GGC CAG CGG CTG ATG AGT ACT GCT GGC CGA GGA C | 34 | 76 | 72 |
| 4 | Q5SDM_ShkADHp-CA-Rec1-DDR_R | ACG GCG CCG GCG ATG CGG | 18 | 81 | 72 |
| 5 | Q5SDM_ShkADHp-CA-Rec1_F | GGA TGA GCC CTA ATA AGA CGA CCG CAT CGC C | 31 | 67 | 68 |
| 6 | Q5SDM_ShkADHp-CA-Rec1_R | TCG GCC ACA CCA TCA GCC | 18 | 71 | 68 |
| 7 | Q5SDM_ShkAD430A_F | GAT CCT GAT GGC CCT GCG AAT GC | 23 | 59 | 62 |
| 8 | Q5SDM_ShkAD430A_R | AGG TCA TAG ACC CCT GCT | 18 | 63 | 62 |
| 9 | ShkAI259V_F | GGC GCG GAC AAC GGC GTT GGG CG | 23 | 75 | 70 |
| 10 | ShkAI259V_R | CGC CCA ACG CCG TTG TCC GCG CC | 23 | 75 | 70 |
| 11 | ShkAI340V_F | CGC GTA GTG GCT TGA CCA GAT AGC CCG AGA A | 31 | 76 | 70 |
| 12 | ShkAI340V_R | TTC TCG GGC TAT CTG GTC AAG CCA CTA CGC G | 31 | 76 | 70 |
| 13 | ShkAI390V_F | CAG CAG CGC GTT GAC CGG ATT GTC CTC GG | 29 | 78 | 70 |
| 14 | ShkAI390V_R | CCG AGG ACA ATC CGG TCA ACG CGC TGC TG | 29 | 78 | 70 |
| 15 | ShkAI405V_F | CGC GGT CGA CGA CGC AGC CTT CGC G | 25 | 77 | 70 |
| 16 | ShkAI405V_R | CGC GAA GGC TGC GTC GTC GAC CGC G | 25 | 77 | 70 |
| 17 | ShkAliDDR_F (11497) | ATT TTC TGC AGG CGG TCA CGT CTA GAG GCG GCG GTA CCG GCG TGA ACG GCG ACA GAT CA | 59 | N/A | N/A |
| 18 | ShkAliDDR_R (11498) | ATT TCT GCA GGC CGA GCT CGC CGC CGG ATC CTT CGG CGT TAA GAC CTC CC | 50 | N/A | N/A |
| 19 | ShkADDRli_F (11499) | ATT TAC GCG TGC GCC GCT GCC GAG CTC GCC GCC GGA TCC TTC GGC GTT AAG ACC TCC C | 58 | N/A | N/A |
| 20 | ShkADDRli_R (11500) | ATT TAC GCG TAT CTA GAG GCG GCG GTA CCG GCG TGA ACG GCG ACA GAT CAA G | 52 | N/A | N/A |

**Supplemental Table 2** | Primers used to obtain the recombinant plasmids (Suppl. Table S1) used in this study. * Tm = calculated melting temperature / ** Exp Ta = annealing temperature used experimentally.

| ID | Name | Description | Boundaries* (aa) | Length** (aa) | Mw** (kDa, monomer) |
|---|---|---|---|---|---|
| 1 | ShkA | Full-length, wild-type ShkA | 1-514 | 534 | 55.5 |
| 2 | $ShkA_{Rec1}$ | ShkA Rec1 single-domain construct | 244-373 | 150 | 15.4 |
| 3 | $ShkA_{Rec2}$ | ShkA Rec2 single-domain construct | 366-514 | 168 | 17.7 |
| 4 | $ShkA_{DHp\text{-}CA\text{-}Rec1\text{-}DDR}$ | ShkA DHp-CA-Rec1 construct, lacking the C-terminal Rec2 domain but including the DDR-motif | 1-379 | 399 | 41.3 |
| 5 | $ShkA_{DHp\text{-}CA\text{-}Rec1}$ or $ShkA_{\Delta Rec2}$ | ShkA DHp-CA-Rec1 construct, lacking the C-terminal Rec1-Rec2 linker and the Rec2 domain | 1-366 | 386 | 40.1 |
| 6 | $ShkA_{I259V}$ | Point mutant I259V, full-length ShkA | 1-514 | 534 | 55.5 |
| 7 | $ShkA_{I340V}$ | Point mutant I340V, full-length ShkA | 1-514 | 534 | 55.5 |
| 8 | $ShkA_{I390V}$ | Point mutant I390V, full-length ShkA | 1-514 | 534 | 55.5 |
| 9 | $ShkA_{I405V}$ | Point mutant I405V, full-length ShkA | 1-514 | 534 | 55.5 |
| 10 | $ShkA_{D430A}$ or $ShkA_{D/A}$ | Point mutant D430A, full-length ShkA | 1-514 | 534 | 55.4 |
| 11 | ShkA | Full-length, wild-type ShkA | 1-514 | 539 | 56.4 |
| 12 | $ShkA_{D369N}$ | Point mutant D369N, full-length ShkA | 1-514 | 539 | 56.4 |
| 13 | $ShkA_{liDDR}$ | ShkA with extended linker, before DDR motif | 1-354-li-355-514 | 605 | 63.0 |
| 14 | $ShkA_{DDRli}$ | ShkA with extended linker, after DDR motif | 1-382-li-383-514 | 605 | 63.0 |

**Supplemental Table 3 |** Recombinant proteins used for *in vitro* (1-10) and *in vivo* (11-14) studies (corresponding plasmids are given in Suppl. Table S1). * Numbering with respect to aminoacid (aa) sequence of full-length, wild-type ShkA / ** Including tags and eventual cleavage sites. Molecular weight (Mw) calculated by ProtParam (ExPASy server, https://web.expasy.org/protparam/).
