## Supplementary Figures for "Hybrid histidine kinase activation by cyclic di-GMP-mediated domain liberation"

Fig S1

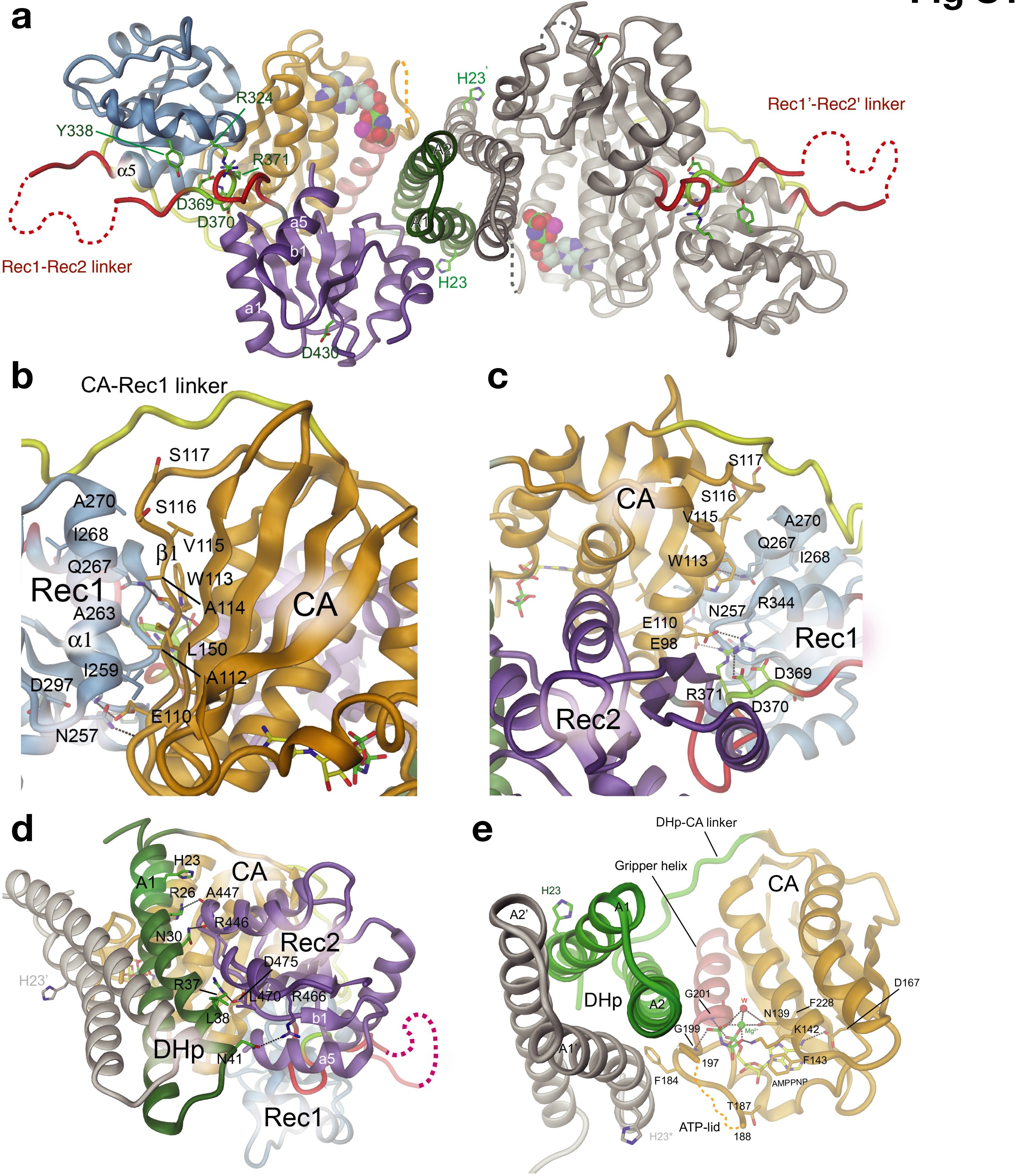


**Supplementary Figure 1. Domain interactions in auto-inhibited, full-length ShkA and close-up view of AMPPNP/ $Mg^{++}$ interactions with ShkA.**

**(a)** ShkA dimer viewed along dyad (as in **Fig. 1b**, bottom), as cartoon representation with exception of Rec1-Rec2 linker that is shown as red thick line with DDR motif in light-green. Residues of the DDR motif, the conserved histidine (H23), and the phospho-acceptor D430 are shown in full. AMPPNP is represented as CPK model.

*Continued on next page.*

**(b, c)** Detailed view of CA/Rec1 and DDR motif/Rec1 interface with interacting residues shown in full. The main (hydrophobic) contact is formed between β2, β4 of CA with α1 of Rec1. There is an extensive hydrophobic interface between these structural elements. D370 from the DDR loop (light-green) makes salt bridges in a network with R344 of Rec1 and D98 of CA (orange).

**(d)** DHp and Rec2 contact with interacting residues shown in full. The hydrophobic residue L38 (helix A1) packs against L470 (Rec2).

**(e)** Cartoon representation of the dimeric DHp bundle and one of the CA domains of full-length ShkA with AMPPNP/$Mg^{++}$ and selected residues shown in full. The phosphate moieties of the ligand are bound to the canonical site composed of the G2 box (G199, G201) and the N-terminal end of the gripper helix (200-213, indian red). The $Mg^{++}$ cation is coordinated by the three phosphates, the side-chain amide of N139 and a water molecule. The adenine base is recognised by the buried D167 and is sandwiched between F143 and F228. The F184 "thumb" ({Bhate et al., 2015}) is found inserted between helices A2 and A1'. The ATP lid is partly disordered.

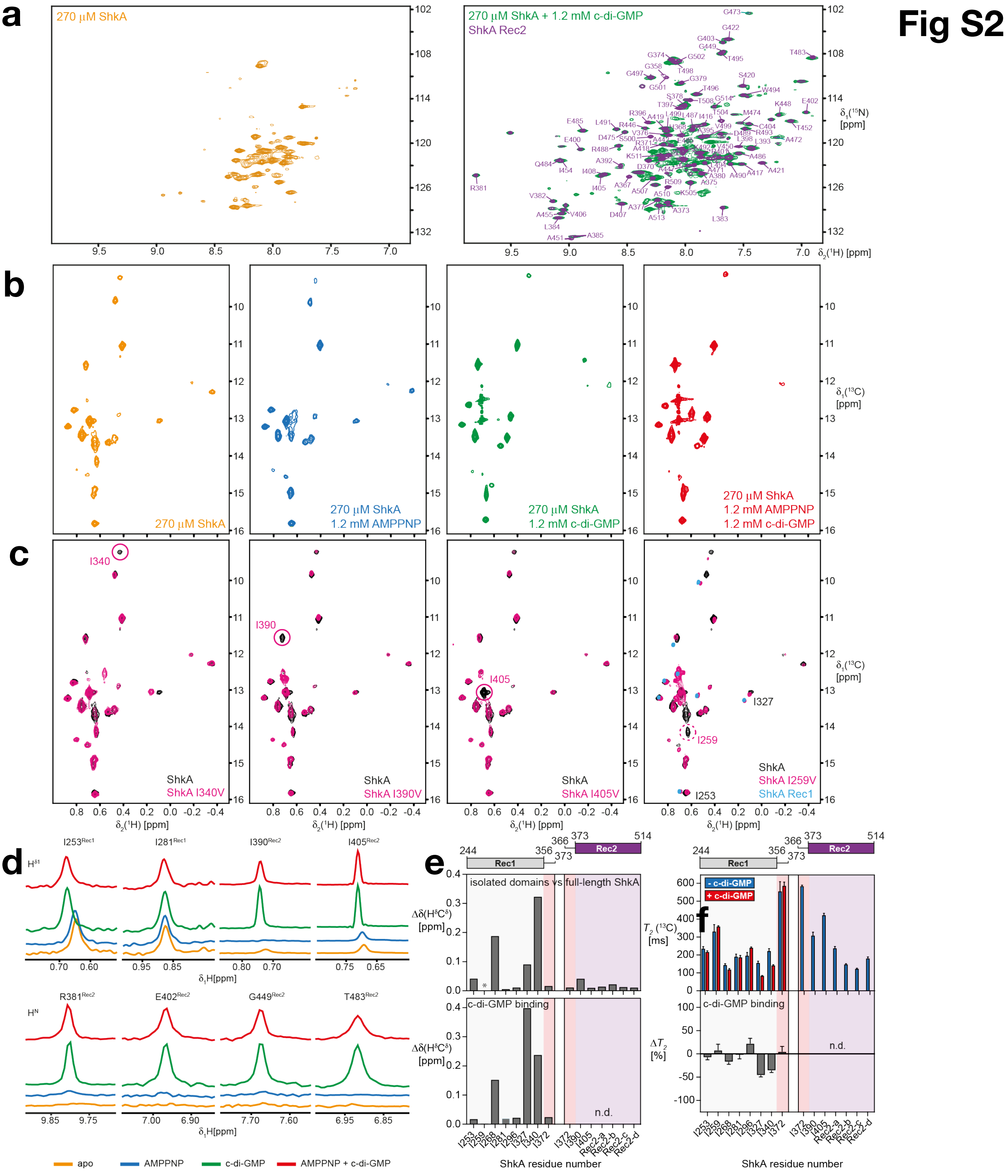


**Supplementary Figure 2. C-di-GMP binding to ShkA leads to release of the Rec2 domain.**

**(a)** 2D [$^{15}$N,$^{1}$H]-TROSY spectra of 270 µM [U-$^{15}$N]ShkA (orange) and overlay of ShkA in its cdG-bound state (green) and the isolated Rec2 domain (purple). Sequence-specific resonance assignments of Rec2 residues are indicated.

*Continued on next page.*

**(b)** 2D [$^{13}$C,$^{1}$H]-HMQC spectra of [$^{13}$C/$^{1}$H$^{\delta1}$-Ile]ShkA in different ligand-bound states, as indicated.

**(c)** 2D [$^{13}$C,$^{1}$H]-HMQC spectral overlays of [$^{13}$C/$^{1}$H$^{\delta1}$-Ile]ShkA wt and single isoleucine mutants, as indicated. In the case of the ShkA I259V mutant also the spectra of the isolated Rec1 domain is shown in the overlay. The resonance assigned the mutated residue is highlighted by a pink circle.

**(d)** Overlay of 1D [$^{1}$H]-cross sections from 2D [$^{15}$N,$^{1}$H]-TROSY and 2D [$^{13}$C,$^{1}$H]-HMQC spectra of representative amide and isoleucine δ1 methyl group protons of ShkA in different ligand-bound states, as indicated.

**(e)** Chemical shift perturbation of isoleucine δ1 methyl groups between the isolated Rec1 and Rec2 domains and full-length ShkA (top panel) and of isolated 225 μM Rec1 upon addition of 0.9 mM cdG (bottom panel).

**(f)** Transverse relaxation times of isoleucine δ1 methyl carbons of isolated Rec1 and Rec2 domains in the absence (blue bars) and presence of cdG (red bars). The bottom panel shows the relative change of $T_2$($^{13}$C) upon addition of cdG ($\Delta T_2$). In e and f, asterisks indicate methyl groups that are either not assigned or line-broadened beyond detection in at least one of the two states.

### Fig S3

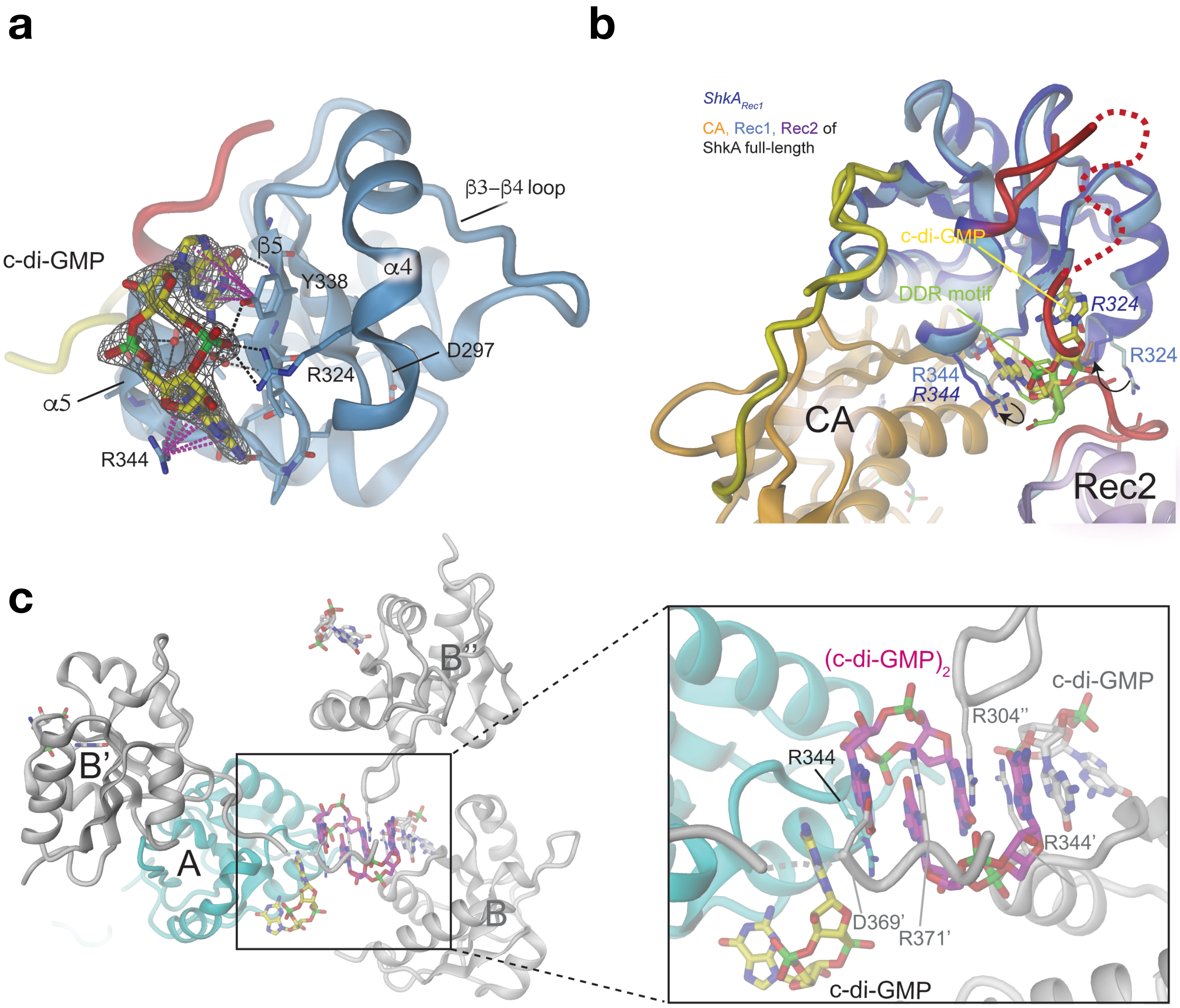


**Supplementary Figure 3. Structural basis of competition between c-di-GMP and DDR for binding to the Rec1 domain of ShkA.**

**(a)** Crystal structure of ShkA$_{Rec1}$ in complex with c-di-GMP shown in full with accompanying Fo-Fc omit map contoured at 1 σ. Monomeric c-di-GMP is bound to the α4 - β5 - α5 face of Rec1. H-bonds are shown by black broken lines and π-cation interactions are indicated by magenta broken. Water molecules are represented as red spheres.

**(b)** Superimposition of ShkA$_{Rec1}$/c-di-GMP onto Rec1 of ShkA full-length. Full-length ShkA domains are distinguished by colour, while the ShkA$_{Rec1}$/c-di-GMP is coloured in dark blue. C-di-GMP and the DDR motif overlap, since they both bind to the same site on the α4 - β5 - α5 face of Rec1. The Rec1 structures are virtually identical with the exception of side-chain orientations of R324 and R344 that appear to adopt to their role in c-di-GMP or DDR motif binding. Thin arrows point from their orientation in the closed (auto-inhibited) full-length structure to their orientation in ShkA$_{Rec1}$.

**(c)** ShkA$_{Rec1}$ crystal packing showing molecule A (cyan) in complex with c-di-GMP (carbons in yellow) linked via a c-di-GMP dimer (carbons in magenta) to the crystallographic symmetry mate B'. The terminal guanine bases of (c-di-GMP)$_2$ form lateral H-bonds with $R_{A344}$ and $R_{B344}$'. In addition, $D_{B369}$' and $R_{B371}$' of the neighboring B molecule are in engaged in lateral H-bonding as well as $R_{B304}$'' of the neighboring B'' molecule.

**Fig S4**

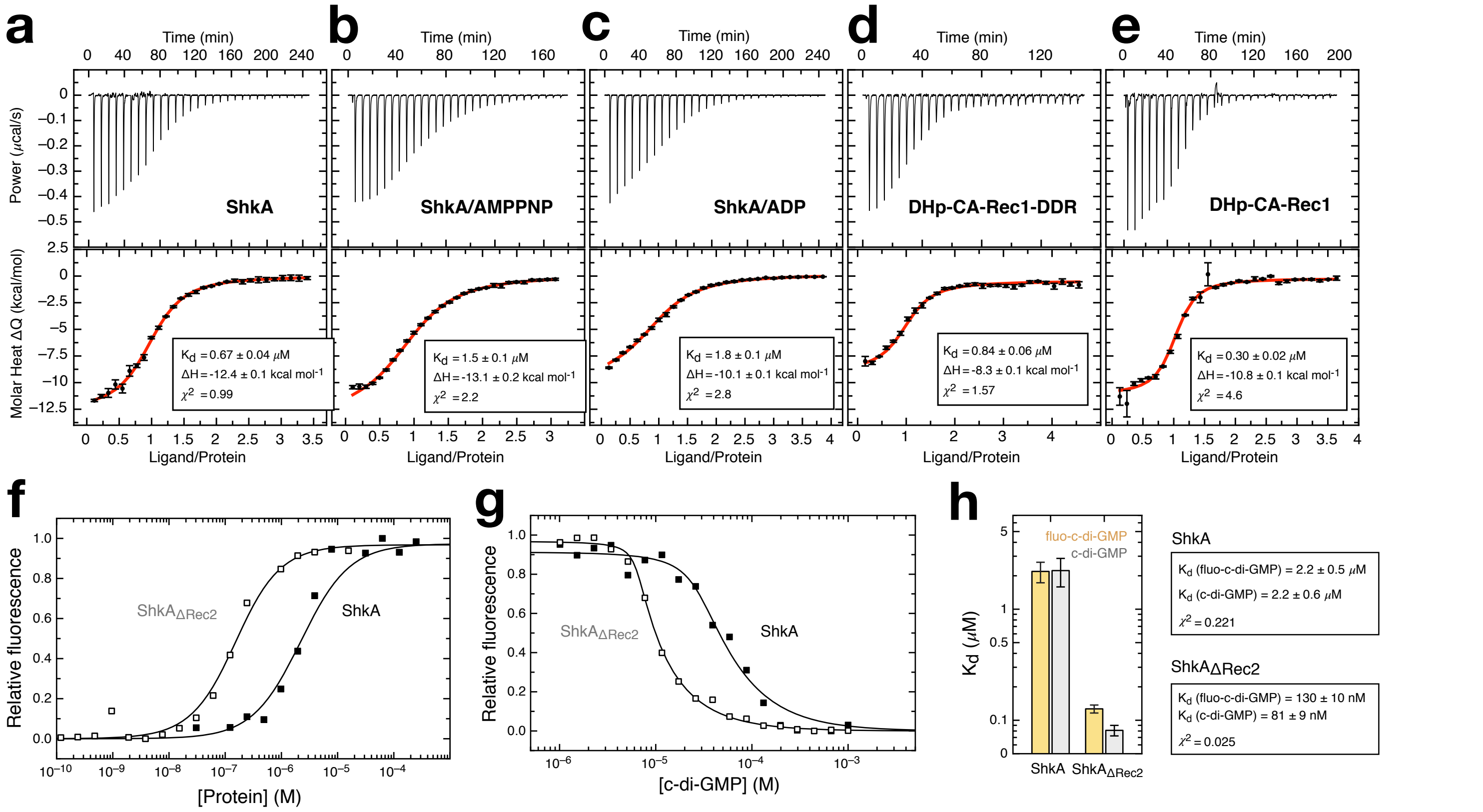


**Supplementary Figure 4: C-di-GMP binding to wild-type ShkA and mutants.**

**(a-e)** Baseline corrected ITC titration data (top) and derived molar heat plots (bottom, with error bars representing peak area errors) of c-di-GMP titration to the indicated ShkA variants. The binding stoichiometry was fixed to N = 1 throughout, given the evidence provided by the crystal structure of $ShkA_{Rec1}$ in complex with c-di-GMP (**Fig. 3a**). Note that for all experiments, the c-di-GMP concentration had to be refined, probably due to the uncertainty of its extinction coefficient. Cell concentration: 10 $\mu$M; syringe concentrations: 0.135 mM (a, b), 0.17 mM (c), 0.2 mM (d), 0.16 mM (e).

**(b, c)** C-di-GMP titration to wild-type ShkA in presence of 50- and 25-fold molar excess of AMPPNP and ADP, respectively.

**(f)** Change in fluorescence intensity upon titration of indicated ShkA variants to 2'fluo-AHC-c-di-GMP (fluo-c-di-GMP, 50 nM).

**(g)** Change in fluorescence intensity upon c-di-GMP titration to indicated ShkA variants (ShkA: 25 $\mu$M; $ShkA_{\Delta Rec2}$: 6.25 $\mu$M) in presence of fluo-c-di-GMP (50 nM).

**(f-g)** Lines represent fit of the ligand competition model {Wang1995} to the data.

**(h)** Binding affinities and statistics as obtained by fitting of the data in panels **f** and **g**.

### Fig S5

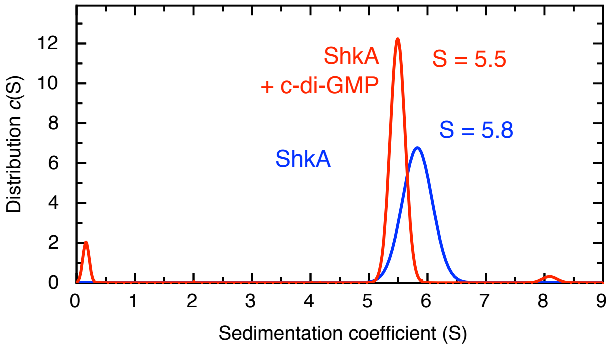


**Supplementary Figure 5: ShkA sedimentation coefficient distribution in absence and in presence of c-di-GMP.**

C-di-GMP binding decreases the sedimentation velocity, which may be attributed to a less compact conformation. [ShkA] = 1 mg/ml = 18 µM, [c-di-GMP] = 45 µM. The distribution was measured by SV-AUC and was fitted by a Gaussian distribution.

**a**

$$E + ATP \underset{k_{-1}}{\overset{k_1}{\rightleftharpoons}} E^* + ADP$$

$$K = \frac{[E^*]\cdot[ADP]}{[E]\cdot[ATP]} = \frac{k_1}{k_{-1}}$$

$$[E] = [E_0] - [E^*]$$

$$[ATP] = [ATP_0] - [ADP]$$

*for* $ShkA_{D/A}$ *and* $ShkA_{\Delta Rec2}$:

$$K = \frac{[ADP]^2}{([E_0]-[ADP])\cdot([ATP_0]-[ADP])}$$

$$[E^*] = [ADP]$$

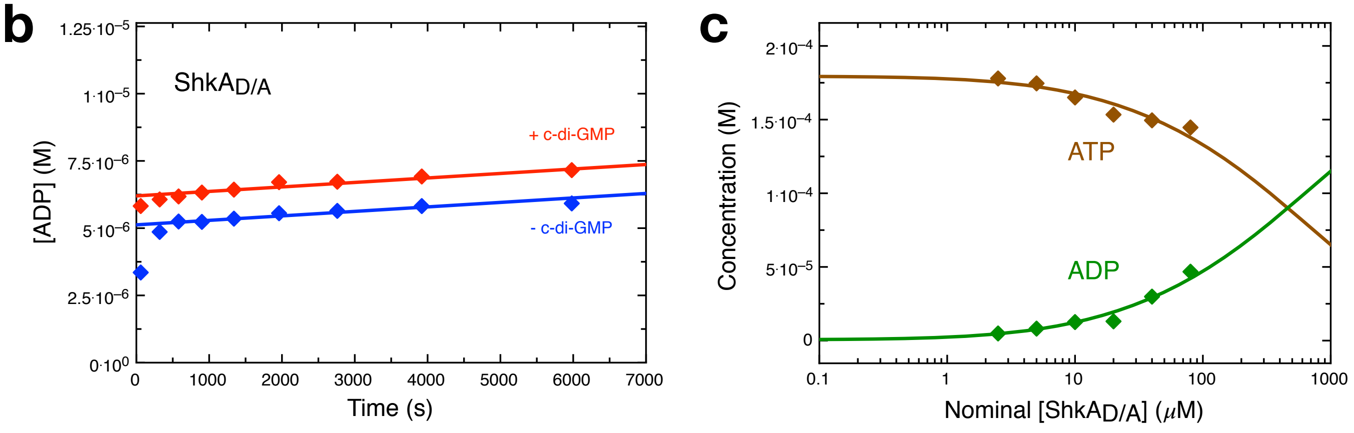


**Supplementary Figure 6. Further characterization of the phosphotransfer deficient $ShkA_{D/A}$ mutant.**

**(a)** Chemical scheme of a reversible autophosphorylation reaction and relationship between equilibration constant K and final (equilibrium) ADP concentration (ADP) for an experiment starting with an initial enzyme concentration $E_0$ and an initial ATP concentration ATP0.

**(b)** Long-term time course of $ShkA_{D/A}$ (10 µM) catalyzed ADP production phosphorylation, compare with Fig. 5d. After the initial fast phase of auto-phosphorylation, there is only a very slow linear increase in the ADP concentration (< $2{\cdot}10^{-5}$ $s^{-1}$) which is attributed to non-specific dephosphorylation of the phosphohistidine followed by re-phosphorylation. The experiment demonstrates that exquisite stability of the phosphorylated histidine at the employed conditions.

**(c)** ADP and ATP concentrations after incubation (15 mins) of $ShkA_{D/A}$ at the indicated concentrations with 80 µM ATP. Data were globally fitted with the reversible (E + ATP <-> E-P + ADP) equilibrium model (panel a, see also Methods) yielding K = 0.11 ± 0.02, similar to the value obtained from the ADP titration experiment (Fig. 5e).

### Fig S7

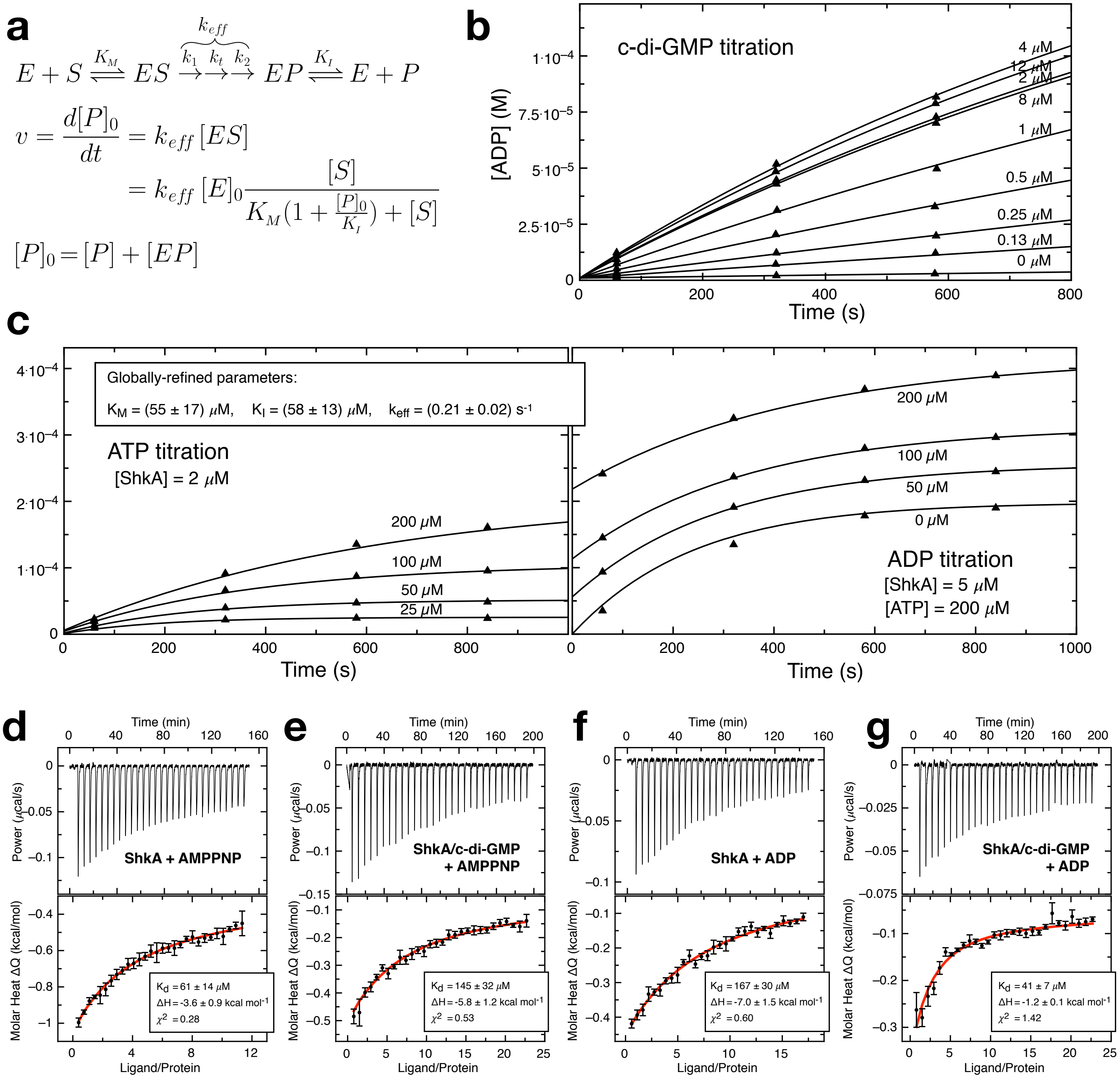


**Supplementary Figure 7. Enzymatic characterization and c-di-GMP affinity of ShkA.**

**(a)** Competitive product-inhibition model based on Michaelis-Menten kinetics. The model was used to obtain the $K_M$, $K_I$ and $k_{cat,eff}$ constants upon fitting the ADP progress curves of panel c.

**(b)** ADP progress curves in presence of the indicated concentrations of c-di-GMP, ATP and ShkA. Data were fitted with the model of panel a to obtain $k_{cat,eff}$ as function of c-di-GMP as plotted in Fig. 5b. The $K_M$, $K_I$ parameters were fixed to the values given in panel c. Scaling factors were fixed to their experimentally obtained values (calibration curves not shown). ATP and ADP concentrations were refined, but deviated maximally by 1% from the nominal concentrations.

*Continued on next page.*

**(c)** ADP progress curves in presence of a 2.5-fold molar excess of c-di-GMP with the indicated starting concentrations of nucleotides (ATP, left; ADP, right) and ShkA. Data were globally fitted to the competitive product-inhibition model shown in panel a and yielded $K_M$ = 54.5 µM, $K_I$ = 58.3 µM and $k_{cat,eff}$ = 0.21 s-1. Scaling factors were fixed to their experimentally obtained values (calibration curves not shown). ATP and ADP concentrations were refined, but deviated maximally by 10% from the nominal concentrations.

**(d - g)** Baseline corrected ITC titration data (top) and derived molar heat plots (bottom) with error bars representing peak area errors. The binding stoichiometry was fixed to N = 1 throughout. Cell concentration: 10 $\mu$M; syringe concentrations: 0.5 mM (d), 0.75 mM (f), 1 mM (e, g).

**(d)** AMPPNP titration to wild-type ShkA.

**(e)** AMPPNP titration to wild-type ShkA in presence of 10-fold molar excess of c-di-GMP.

**(f)** ADP titration to wild-type ShkA.

**(g)** ADP titration to wild-type ShkA in presence of 10-fold molar excess of c-di-GMP.
